# Characterization of bacteria colonizing the mucosal layer of the gastrointestinal tract of Atlantic salmon farmed in a warm water region

**DOI:** 10.1101/2024.11.23.625004

**Authors:** Kate Hall, Chantelle E Reid, Josephine Hamlett, Damanpreet Kaur, Qi Zhi Tan, Richard S Taylor, Andrew Bissett, Barbara F Nowak, John P Bowman

## Abstract

Atlantic salmon (*Salmo salar*) farmed in seawater in Tasmania (*lutruwita*) can experience temperatures close to their thermotolerance limit during summer. Gut microbiome data from eight successive annual surveys and a specific survey of GI tract mucosa and digesta bacterial cross-sectional distributions indicated members of the genus *Aliivibrio*, *Vibrio*, and an unclassified Mycoplasmoidaceae are the main colonizers of the gut mucosal layer in Tasmanian farmed salmon. Peak abundance levels are reached after 7-8 months, corresponding to the late summer, following the transfer of hatchery smolt to seawater farms. Salmon *Aliivibrio* isolates comprise three novel non-bioluminescent species distinct from species found in the northern hemisphere growing regions. Along with other *Aliivibrio* species these species share many genes required for host colonization and biofilm formation and also include species and strain-level dependent features. Two of the novel *Aliivibrio* species surprisingly possessed genes for cytolethal distending toxin while the more predominant species lacked any known virulence genes. The overall observations suggest a small number of species actively colonize the mucosal layer of Atlantic salmon farmed in Tasmania and that this process is strongly influenced by the environmental temperature.

## Introduction

In Tasmania (Australia), also referred to as *lutruwita*, the Atlantic salmon industry is a major economic activity and relies on sea cage farming. The industry operates at the high end of the temperature scale owing to summer sea surface temperatures being on average 15-20°C while Atlantic salmon grow optimally at 13-16°C (Calado et al., 2021). There are concerns that sea surface temperatures in the south-east of Tasmania will increase in the longer term owing to the ongoing topicalization of the Eastern Australian Current (Oliver et al., 2017). An unprecedented 117-day “heat wave” during the summer of 2015-2016, where surface (5 m depth) temperatures reached almost 23°C caused obvious negative effects on local farmed salmon health and downstream marketability (Wade et al., 2019). It is predicted increased warming will have potentially detrimental effects (Grünenwald et al., 2019; Hudson et al., 2022) on the Atlantic salmon industry in some regions (Meng et al., 2022).

Gut microbiome studies on Atlantic salmon and other related marine farmed finfish have been performed under numerous contexts. This mainly has been done to understand the effects of environmental conditions, comparing life stages, establishing health status indicators, and connecting this data to farm performance and feed formulations (Egerton et al., 2018). One of the challenges in such studies has been understanding what the main players in the microbiome are, how this compares between growing regions, and what the microbiome means for Atlantic salmon health and performance. The overall commercial system is relatively complex and only approximates the Atlantic salmon anadromous lifestyle. Typically, juvenile salmon are reared in freshwater hatcheries up to the smolt life stage (Li et al., 2023). By increasing light exposure and temperature, maturation and associated osmoregulation (Ytrestøyl et al., 2023) is stimulated mimicking the natural spring-summer transition in physiology of wild salmon. This artificial stimulus enables the smolt to rapidly adapt to seawater conditions, whereafter they are reared in marine farms (Hvas et al., 2021). Across this process gut microbiota has been observed to change unidirectionally (Zarkasi et al., 2014; Lorgen-Ritchie et al., 2021; Wang et al., 2021). The process of change has been associated with fish age, feeding rates, and weight gain (Zhao e al., 2020).

Studies to date clearly indicate that bacteria are predominant and establish populations in the gastrointestinal mucosal layer and the digesta of Atlantic salmon. The main autochthonous adherent taxa, distribution within the gut (pyloric caeca to distal gut), and aquaculture region differences are now being established at high resolution (Vera Ponce de Léon et al., 2024). A major autochthonous species present in wild Atlantic salmon has been described as “Candidatus Mycoplasma salmoninarum” (Rasmussen et al., 2023), a member of genus *Malacoplasma* in the family Mycoplasmoidaceae. This species is found in abundance in the gut of wild post-smolt (Llewellyn et al., 2016) as well as in salmon farmed in sea pens world-wide. Abundance ranges from being nearly completely dominant to only a sporadic presence (Zarkasi et al., 2014; Dehler et al., 2017; Fogarty et al., 2019; Bozzi et al., 2021; Huyben et al., 2020; Wang et al., 2021; Brealey et al., 2022). In the Tasmanian salmon growing region (147°E 43°S) Vibrionaceae (*Aliivibrio*, *Vibrio*, *Photobacterium*) occur at high levels in adult fish (Reid et al., 2024). Vibrionaceae are also common in ocean-dwelling wild salmon and other farming regions (Wang et al., 2020, 2021, 2023; Lorgen-Ritchie et al., 2021, Llewellyn et al., 2016; Brealey et al., 2022; Huyben et al., 2020; Godoy et al., 2015; Dhanasiri et al., 2023; Vera Ponce de Leon et al., 2024). Similar colonization patterns also occur in other anadromous, farmed salmonids such as rainbow trout (Rimoldi et al., 2021) and chinook salmon (Ciric et al., 2019; Zhao et al., 2020; Ziab et al., 2023). For example, in chinook salmon, farmed extensively in New Zealand, *Aliivibrio* and *Photobacterium* predominate in fish in marine farms while *Aliivibrio* does not occur in fish reared in alpine freshwater systems (Zhao et al., 2020).

Certain members of Vibrionaceae cause disease in Atlantic salmon and the most problematic have been managed by vaccination (Skåne et al., 2022). More avirulent species may be predominant in fish which produce signs of dysbiosis, including reduced voluntary feeding and expression of fecal casts. Reduced feeding can lead to anorexic-like states in a proportion of fish (Hevrøy et al., 2012). Casts represent sloughed intestinal mucosa appearing as white to yellow fecal matter (Reid et al., 2024). Both phenomena occur in fish that are thermally stressed though whether Vibrionacaeae are directly involved in dysbiotic symptoms is yet to be shown. *Aliivibrio* are of particular interest not just due to their potential to cause disease but also possess a capacity to act as probionts (Klakegg et al., 2020). These characteristics suggest an inherent capacity for host colonization. Significant knowledge of host interactions obtained for the model species *Aliivibrio fischeri* (Visick et al., 2021) provides a foundation for functional comparison of different *Aliivibrio* species colonizing fish GI tracts.

Due to the predominance of Vibrionaceae in marine farmed fish, there is usefulness in understanding their biology. This includes understanding capabilities related to persistent colonization, growth *in vivo,* response to water temperatures, and connectivity to dybiosis. In this study we examined the colonization of Atlantic salmon reared in marine pens located in the D’Entrecasteaux Channel region of Tasmania. Unlike previous studies we performed an analysis of gut microbiome samples collected over multiple years (2010 to 2018) to determine colonization patterns. Secondly, we compared the distribution of bacteria through the GI tract comparing different gut sections in sets of fish collected from two separate cohorts in summer and winter. This was done to answer the question more definitively of what bacteria show proliferation in the gut mucosa versus simply passive transit via feed or water. To try to understand the traits of active colonizing bacteria in the gut of Atlantic salmon we also isolated strains from salmon digesta samples and sequenced their genomes. From this we examined their taxonomy and relevant genes the strains possess that enable host colonization. Finally, we also wanted to know if gut colonizers possess virulence genes maintaining a focus on the relevance of strain colonization as a facet of the health, welfare and farm performance of Atlantic salmon, especially in commercial settings. From these studies we can show Atlantic salmon is colonized actively by a number of distinct species with strong capability for growth in the Atlantic salmon GI tract mucosal layer.

## Materials and Methods

### Sea pen sampling

For sea pen derived fish, data in this study included several different surveys performed between 2010 and 2018 in the southeast of Tasmania (Table 1, Figure S1). Four of these surveys have been previously published where the purpose of the research was to understand aspects of aquaculture different to what is being explored in this study. These previous studies included studies on the effect of diet formulations, fecal consistency, short-term sampling after feeding, and interactions of salmon with Australian fur seals (Zarkasi et al., 2014, 2016; D’Agnese et al., 2023; Reid et al., 2024). New data are from unpublished surveys of farmed Atlantic salmon in the southeast of Tasmania (Table 1). In each studied cohort, smolt had been put to sea over winter (July to August). Samples were collected between 0 to 3 weeks prior to transfer from two hatcheries and then from 2 to 56 weeks after smolt transfer. Digesta samples were obtained by fecal stripping as detailed below. Data from Reid et al. (2024) was also used to investigate gut mucosa and digesta samples of fish that had been euthanized and dissected, Methods for that sampling is described in detail in that paper.

**Table 1.** Experimental datasets utilized in the study including design, data sources and related information.

| Experiment-type | Location of sampling | 16S rRNA primers | Sequencing method | Time period samples collected | Number of samples per time point | Fish weigh range (kg) | Reads per sample: average (range) | Reference | NCBI sequence read accession code | Animal ethics approval no. |
| --- | --- | --- | --- | --- | --- | --- | --- | --- | --- | --- |
| Farm-based | Meads Creek, Port Esperance (1) | V3-V5 | Pyroseq-uencing | Aug 2010 – Apr 2011 | 20, 49, 50, 50, 50 | 0.4-4.1 | 3900 (654-21356) | Zarkasi et al. 2016 | PRJNA1021267 | DPIPWE 30/2009-10 |
| Farm-based | Robert's Point lease, D'Entrecasteaux Channel (2) | V3-V5 | Pyroseq-uencing | Jul 2011 – Aug 2012 | 6, 12, 12, 11, 12, 11, 10, 12, 12 | 0.5-1.8 | 3994 (489-13691) | Zarkasi et al. 2014 | PRJNA1021267 | UTAS A0012001 |
| Farm-based | North West Bay (3) | V3-V5 | Illumina MiSeq | Nov 2013 – Jun 2014 | 9, 8, 11 | Not available | 146271 (22193-404791) | This study | PRJNA1021367 | UTAS A0013971 |
| Farm-based | Red Cliffs, D'Entrecasteaux Channel (4) | V1-V3 | Illumina MiSeq | Mar 2014 – Jun 2014 | 64, 49, 76 | 1.2-2.2 | 221637 (6932-627716) | D'Agnese et al. 2023 | PRJNA979964 | UTAS A0013971 |
| Farm-based | North West Bay (3) | V1-V3 | Illumina MiSeq | Jan 2015 – Apr 2015 | 22, 29, 18 | Not available | 55446 (1240-118582) | This study | PRJNA1021684 | UTAS A0013971 |
| Farm-based | Soldier's Point, D'Entrecasteaux Channel (5) | V3-V5 | Illumina MiSeq | Dec 2015 – May 2016 | 16, 30, 21, 11 | Not available<br>Not available | 31740 (1254-81741) | This study | PRJNA1021687 | UTAS A0015452 |
| Farm-based | Red Cliffs lease, Sheppards lease, D'Entrecasteaux Channel (4, 6) | V1-V3 | Illumina MiSeq | Jan 2018, Jul 2018<sup>a</sup> | 182, 196 | 2.1-2.3 | 13820 (602-51163) | Reid et al. 2024 | PRJNA1025921, PRJNA1026149 | UTAS A0016307 A0016588, A0017010 |
| Hatcheriesf arms | feed pellets, water | V1-V3 | Illumina MiSeq | Jan 2018 – Jul 2018 | 41, 38 | Not applicable | 13727 (6999-31697), 13350 (2924-51553) | This study | PRJNA1025868 | Not applicable |
| Hatcheriesf arms | Robert's Point Lease, Sheppards leases, D'Entrecasteaux Channel (4, 6) | V1-V3 | Illumina MiSeq | Jul 2017- Jun 2018 | 234 | 0.06-3.9 | 23496 (4220-53964) | This study | PRJNA1025921, PRJNA1026149 | UTAS A0016307 A0016588, A0017010 |

<sup>a</sup>Samples were collected from fore-gut, mid-gut, and hind-gut digesta and mucosa as well as by faecal stripping in the 24 h period after fish fed (Reid et al. 2024).

### Digesta, feed pellet and water sampling

Groups of Atlantic salmon were captured using a large seine net and the net was shallowed to crowd the fish to ensure that they were randomly mixed. Fish were sampled randomly by using a dip-net. Captured fish were transferred into a 300 L aerated tub containing 17 ppm Aqui-S. For faecal stripping, ventral surfaces of anaesthetized fish were wiped with ethanol to minimize contamination, and samples were obtained by gently pinching and massaging the fish from the midline above the pelvic fins down towards the vent. Initial samples stripped from each fish was visually scored using the faecal consistency scoring method of Zarkasi et al., (2016). Following faecal scoring 200 to 300 mg of digesta was aseptically collected and placed into a sterile 2 ml tube. The samples were then snap frozen in a dewar containing liquid nitrogen and transported back to the laboratory and stored at – 80 °C until DNA was extracted. All sampling from either hatcheries or farms was performed under University of Tasmania Animal Ethic Permits as listed in Table 1. Considerable steps were taken to minimize cross contamination by using sterile surgical sheets and by disinfecting dissecting equipment by immersion into 95 % (v/v) ethanol and then a 10 % (v/v) sodium hypochlorite solution for DNA decontamination (Nilsson et al., 2022). Water samples were also collected at all sampling events by submerging and filling autoclaved 2 L glass laboratory bottles from either inlet water or directly from tanks as a representation of microbes in the surrounding environment. This water was then filtered through a 0.22 µm Sterivex (Merck Millipore) filter units using a peristaltic pump, and the filters were frozen at −80 °C for further processing. The pump was sterilized between samples by pumping through 0.1% NaClO, flushing through and remaining NaClO solution residual was rinsed away using sterile distilled water. Due to possible influence of DNA from feed pellets (Karlsen et al., 2022), feed pellet samples were collected from automatic feeders by transferring directly to sterile zip lock bags. All samples were kept frozen at −80 °C until processed.

### DNA extraction of samples and sequencing

Total genomic DNA from digesta and feed pellets (0.25 g frozen weights) was extracted following manufacturer’s specifications using the QIAamp DNA Stool Kit (Qiagen Cat. No. 51604). To ensure full bacterial lysis, the temperature for lysis incubation temperature was increased to 95 °C and incubation time was increased from 10 min for all samples. Isolation of genomic DNA from Sterivex filtered water samples were processed using the DNeasy PowerWater Sterivex Kit (Qiagen Cat no. 14600-50-NF) following manufacturer’s specifications. The concentration of DNA extracted was determined using a NanoDrop 8000 spectrophotometer (Thermo Fisher Scientific).

### Sequence processing and classification

Sequencing was performed at the Ramaciotti Centre for Genomics, University of NSW, Sydney, Australia using the Illumina MiSeq platform. For this 16S rRNA genes were amplified using V1/27F 5’-AGA GTT TGA TYM TGG CTC AG-3’; (Lane 1991) and V3/519R 5’-GWA TTA CCG CGG CKG CTG-3’ (Turner et al., 1999) primers. The Illumina pipeline used automatically trimmed adaptors from the 3’ prime end of sequences. Paired end reads were joined in SEED2 (Větrovský et al., 2018) using FastqJoin 1.1.2. The joined files were then quality filtered in SEED2 (using Mothur 1.34.4) without pre-clustering, yielding reads with mean median quality scores of ≥35 (range 35 to 39). FASTA files of the filtered reads were generated and then processed by VSEARCH (Rognes et al., 2016) to remove chimeric sequences and to create a list of reads clustered at 1% dissimilarity. The output cluster FASTA files were then combined into a single FASTA file in SEED2, and the process of clustering and chimera removal was repeated to create a single non-redundant OTU dataset covering all the samples. The OTU list was then classified to the genus level using megaBLAST v. 2.2.26+ against the Silva 138.1 non-redundant 16S rRNA gene database (Quast et al., 2013) in SEED2. (Supplementary datafile 1).

### Enumeration, isolation and 16S rRNA gene analysis of salmon gut bacteria

Isolation and enumeration of bacteria from Atlantic salmon digesta utilized marine agar (MA - 0.5% w/v Bacto-peptone, 0.2% w/v yeast extract, 3.5% w/v seawater salts, 1.5% w/v agar), Thiosulfate Citrate Bile Sucrose agar (TCBS, Oxoid), and de Man-Rogosa-Sharpe agar (MRS, Oxoid). For these experiments fixed wet weight amounts of digesta samples (typically 0.2 to 1.0 g) were suspended and serially diluted in 3% (w/v) sterile seawater salt solutions, spread onto the surface of MA, TCBS and MRS and then incubated at 25°C for up to 7 d. Counts were calculated as CFU per g of feces (wet weight). For various salmon samples (mainly obtained between 2015/2016 to 2017/2018) bacterial isolates were obtained from MA plates. Individual colonies were picked from agar plates and purified on MA. Pure cultures were cryopreserved in marine broth that included 30% (v/v) glycerol and stored at −80°C. DNA was extracted from each strain using the Ultraclean Microbial DNA Extraction kit (Qiagen). Isolates were identified using 16S rRNA sequence gene analysis. Amplification of 16S rRNA gene used 10 pmol/μl 27F and 1492R 5’-GGT TAC CTT GTT ACG ACT T-3’ (Lane, 1991) primers. PCR utilised MyTaq PCR reagent (Bioline) and included 1 μl (20-50 ng) of template DNA. Thermocycler (Peltier PTC200, MJ Research) conditions used were 95°C 1 min (1 cycle); 95°C – 30 s, 50°C – 30 s, 72°C – 30 s (34 cycles); 72°C – 10 min (1 cycle); 10°C soak. PCR products were purified and then sequenced using BigDye chemistry using an AB3730xl sequencer instrument (Applied BioSytems). Sequence end regions were trimmed to remove poorly resolved base calls and aligned to reference sequences downloaded from the NCBI database. Phylogenetic trees were generated using the BioNJ algorithm in NGphylogeny.fr (Lemoine et al., 2019) and visualized using the Interactive Tree of Life (Letunic & Bork 2021).

### Growth analyses and biofilm formation ability

To assess responses to media salinity the cultures were grown in marine broth in which sea salts was replaced by different concentrations of NaCl (0 to 10% NaCl). Bile salt tolerance was assessed similarly with marine broth supplemented with bile salts no. 3 (0 to 5%, Oxoid). The ability of isolates to form biofilms was assessed using the crystal violet assay in 96 well polystyrene trays and on acid-washed glass surfaces as previously described (Hussa et al., 2008, Hansen et al., 2014). The motility of cultures (incubated in marine broth for 40 to 72 h at 20°C) was tested under phase contrast microscopy using wet mounts. The biokinetic growth properties of the salmon isolates was assessed using a gradient temperature incubator (Mellefont & Ross 2003). Representative isolates were grown in marine broth and marine broth supplemented with 1% (w/v) ox bile salts at 25°C for 48 h. Inoculum was transferred to a fresh media and grown again for at 25°C for 24 h. Cultures were diluted to 10^4^ to 10^5^ CFU/ml in fresh marine broth that had been aliquoted into L-shaped 15 ml test tubes trays. The L-shaped tubes were placed into a custom-built aluminum block temperature gradient incubator (Terratec, Kingston, Tasmania) and incubated at temperatures ranging from 2°C to 40°C. Optical density readings were taken regularly using a Spectronic 200 (ThermoScientific) until no change in absorbance values was evident.

### Genome assembly and annotation

Genome sequence data were obtained using either the Illumina HiSeq or the NovaSeq 6000 platforms in which 200 or 250 bp pair-ended sequences were generated. Fastq files were uploaded onto the Galaxy Australia cloud server and assemblies generated using Unicycler v. 0.050 and samtools v. 1.15.1 (Wick et al., 2016). The default parameters were used for the assembly process. Genomes were then annotated using the PANNZER2 server (Törönen & Holm, 2022). Non-coding RNA genes were detected using BARRNAP v. 0.9 (Torsten Seemann, https://github.com/tseemann/barrnap) as implemented in Galaxy.

### Genome-based taxonomy

Genomes were compared using ortho-ANI (Lee et al., 2016) and GGDC 3.0 model 2 (Meier-Kolthoff et al., 2013) to discern putative species level groups assuming 95% and 70% demarcate approximate boundaries for species, respectively. Similarity values from both approaches were used to create dendrograms in Primer 7 (Primer-E, Auckland, New Zealand) using the complete linkage method. The 16S rRNA genes directly amplified from isolates and those annotated in genomes were compared including determination of variants from the genome data using BARRNAP v. 0.9. The V1 to V3 regions of sequences from isolates and available genomes of other *Aliivibrio* species, as defined by Klementsen and colleagues (2021), were compared in a tree created using the BioNJ algorithm as described above.

### Compositional and differential abundance analysis

Comparisons of relative abundances of bacteria taxa enriched in digesta and mucosal samples in different parts of the gastrointestinal tract of Atlantic salmon. For this the microbiome of the hind gut and mid-gut digesta and mucosal samples were compared with the foregut samples. In this analysis it was assumed proliferating bacteria would become more abundant progressing from the fore-gut through to the hind-gut. Similarly, it was assumed feed derived DNA would undergo dilution. The analysis started with read abundances of individual OTUs transformed to CLR values (Aitchinson 1982). These values allowed a paired-analysis approach in which the difference in CLR values (ΔCLR) were calculated to estimate the relative abundances of OTUs between gut section samples. Samples collected from fish during both summer and winter were analysed. This resulted in four ΔCLR datasets possessing 27 paired samples for each OTU analysed. The Wilcoxon signed rank test was used to compare OTUs in the paired samples. Assessment of the data indicated that as the OTU abundance became small data asymmetry tended to increase appearing as small negative numbers. To check the Wilcoxon data the Mann Whitney U test was also performed to help eliminate false positives. We surmised from the results that for meaningful differences in OTU abundances between gut section samples to occur ΔCLR values had to be large. False discovery rate (FDR) testing of the Wilcoxon (CLR) p-values, using the method of Benjamini and Hochberg (1995), indicated OTUs with nominal significance (p <0.05) was achieved with paired ΔCLR values of ≥0.4 and ≤-0.3. These values corresponded to large effect sizes (*r* ≥ 0.5) (Fritz et al., 2012). Thus, by requiring effect size (*r* >0.5) and ΔCLR (>0.4 or <-0.3) to be large avoided false positives. OTUs with ΔCLR that had passed these criteria had significant W-values (p<0.05, FDR) for at least one of the four dataset comparisons. These OTUs tended to be both abundant and prevalent in digesta and mucosa samples. The resultant OTU ΔCLR values were arrayed into a heat map. The overall paired dataset results and OTU ΔCLR values across the paired datasets was hierarchically clustered utilizing complete linkage of un-centred correlations. This was completed in Cluster 3 (de Hoon et al., 2004). The heat map was created using TreeView 3.0 (Saldanha 2004).

### Colonization and virulence-based features annotated from genome data

Genes associated with host colonization associated phenotypes, including adherence, biofilm formation, quorum sensing, interbacterial competition and host-directed virulence were surveyed in *Aliivibrio* as well as *Vibrio scophthalmi* genomes. This survey was based on mechanistic studies of gene and protein function performed to date in mostly *A. fischeri* and *V. cholerae*, coupled to data gleaned from the online databases KEGG (Kanehisa & Goto 2000) and VFDB (Liu et al., 2022). The presence of target proteins for this survey was assessed using protein comparisons using BLAST-P in NCBI with confirmation sequences matched on the basis of similarity (>40%), length (>90%) and conserved domain structure. NCBI database contains 5 available assembled *V. scophthalmi* genome sequences which were used in these analyses.

## Results

### Multiyear surveys of gut microbial profiles and proliferating taxa

Gut microbiome taxonomic profiles (supplementary datafile 1) from Atlantic salmon digesta and mucosa were collected between 2010-2018 (Table 1). This dataset was used to determine which bacterial taxa consistently predominate each year and whether these taxa show evidence of proliferation over time. The sampling covered fish reared from smolt through to adults (up to 56 weeks at sea) across six different locations within the D’Entrecasteaux Channel region of Tasmania (Figure S1). Smolt are transferred from hatcheries to farms in winter when water temperatures typically are 10-13°C. Summer water temperatures in the D’Entrecasteaux Channel range from 15 to 20°C. Proliferating taxa were defined as those exhibiting an increasing proportion of reads over time after smolt had been transferred to farms, are present in a high proportion of samples across the surveys, and eventually occur at a high relative abundance given time. The profiles of temporally categorized samples are shown in Figure S2. The OTUs fulfilling the criteria for proliferation the best included those classified as *Aliivibrio*, *Vibri*o, and *Photobacterium* and unclassified *Mycoplasmoidaceae*. Collectively, these taxa made up 78% of total reads (of 4.0 × 10^7^ total reads) and were highly prevalent in samples (91-97% overall).

The abundance of these genera is shown in relation to the time fish had spent in marine farms amalgamating data from all surveys (Figure 1). On an individual fish level, there is a high degree of variability, however a clear pattern of increasing CLR abundance over time occurs. To assess how much change, assumed to be due to mainly cell division, occurs over the survey period the averages of the CLR values of the last three timepoints equivalent to the second winter (months 10 to 13) were subtracted from the average of levels in fish sampled in the first winter at sea (months 0 to 2). This increase was estimated to be 3.2 (± 0.3), 2.2 (± 0.7), 2.0 (± 0.8) and 1.0 (± 0.9) log units for *Aliivibrio*, *Photobacterium*, *Vibrio* and unclassified Mycoplasmoidaceae, respectively (Supplementary datafile 2). A number of other taxa were found to partly meet the criteria but were much less abundant and the difference over the annual period, distribution and prevalence was smaller and less consistent. These taxa included *Propionigenium*, *Shewanella*, *Moritella*, *Myroides*, “Candidatus Pelagibacter”, unclassified *Endozoicomondaceae*, unclassified *Mycoplasmatales*, and unclassified Saccharibacteria. The remaining taxa exhibited no evidence of significant changes or show significant decreased abundances (Supplementary datafile 2).

**Figure 1.**
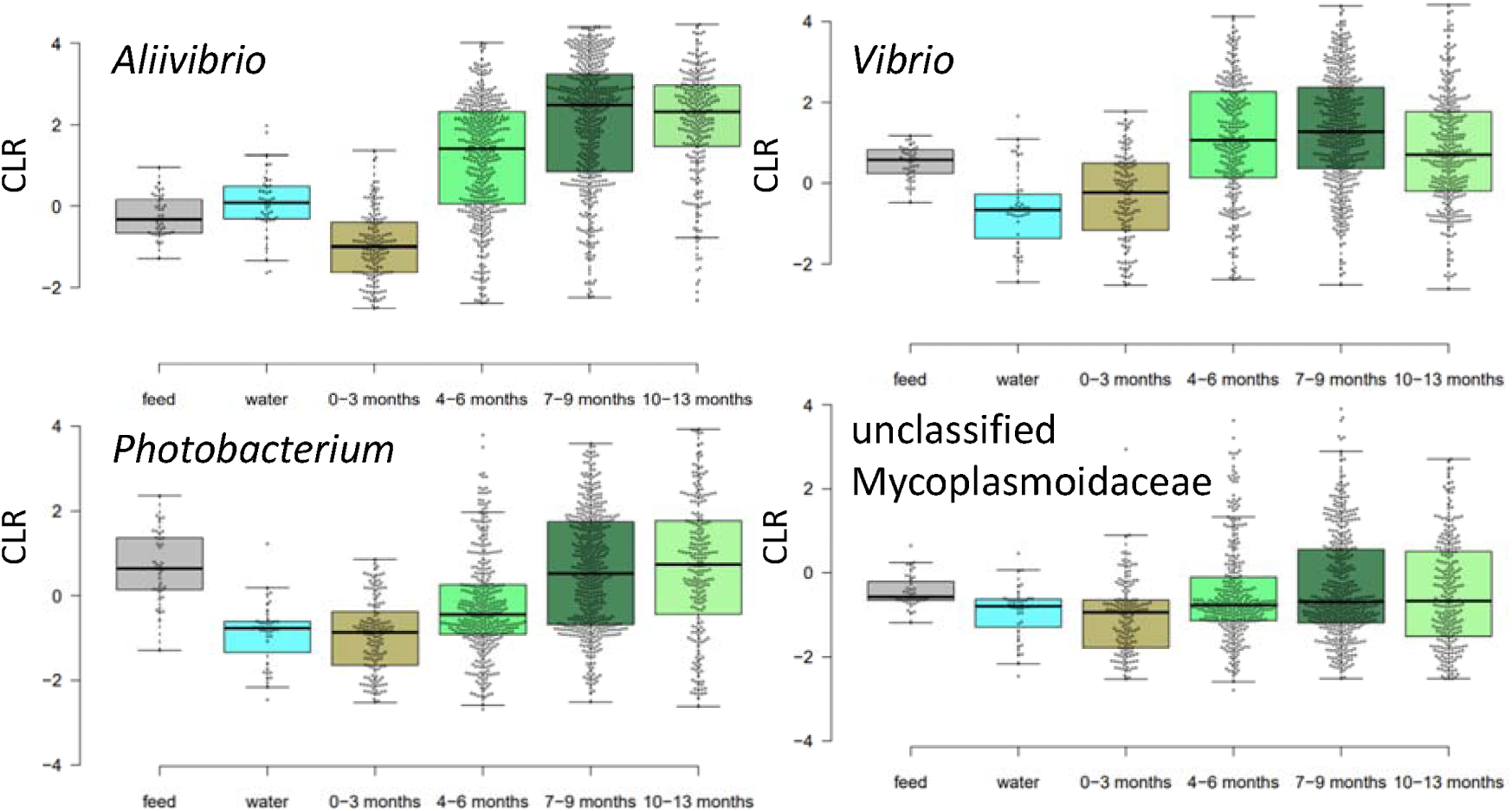
Changes in abundance (CLR) of predominant Atlantic salmon gut microbiome members after being transferred from freshwater hatchery to marine farms in Tasmania.

Based on this data from a low starting abundance at the point of smolt input in winter (July-August) there is an increase in abundance of Vibrionaceae in the spring (October-November) to a peak in the late summer period (March) after which populations begin to plateau amounting to a period of approximately 8 months. *Aliivibrio* had the most definable growth-like sigmoidal response with average CLR values reaching a maximal level by March-April (8 months after transfer) (Figure 2). The response of the unclassified Mycoplasmoidaceae was more muted due to its more sporadic abundance in specimens. This taxon did not appear to show clear evidence of increase until the second autumn-winter period (10-12 months after transfer) had commenced (Figure 2).

**Figure 2.**
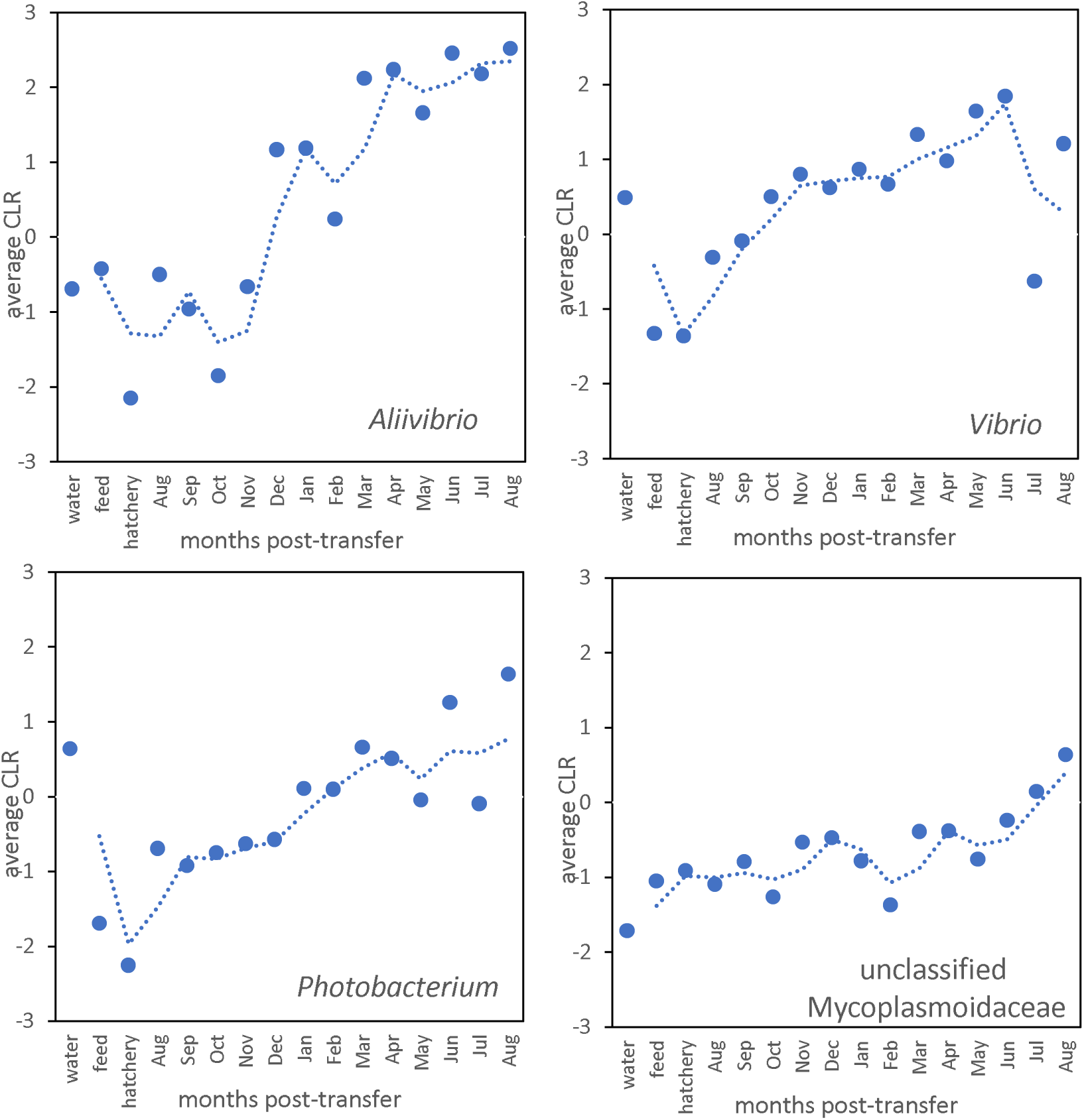
Running average of CLR values integrating all survey data for the most predominant taxa associated with Atlantic salmon gut microbiome. The abundance values are compared to feed, water and hatchery smolt.

**Figure 3.**
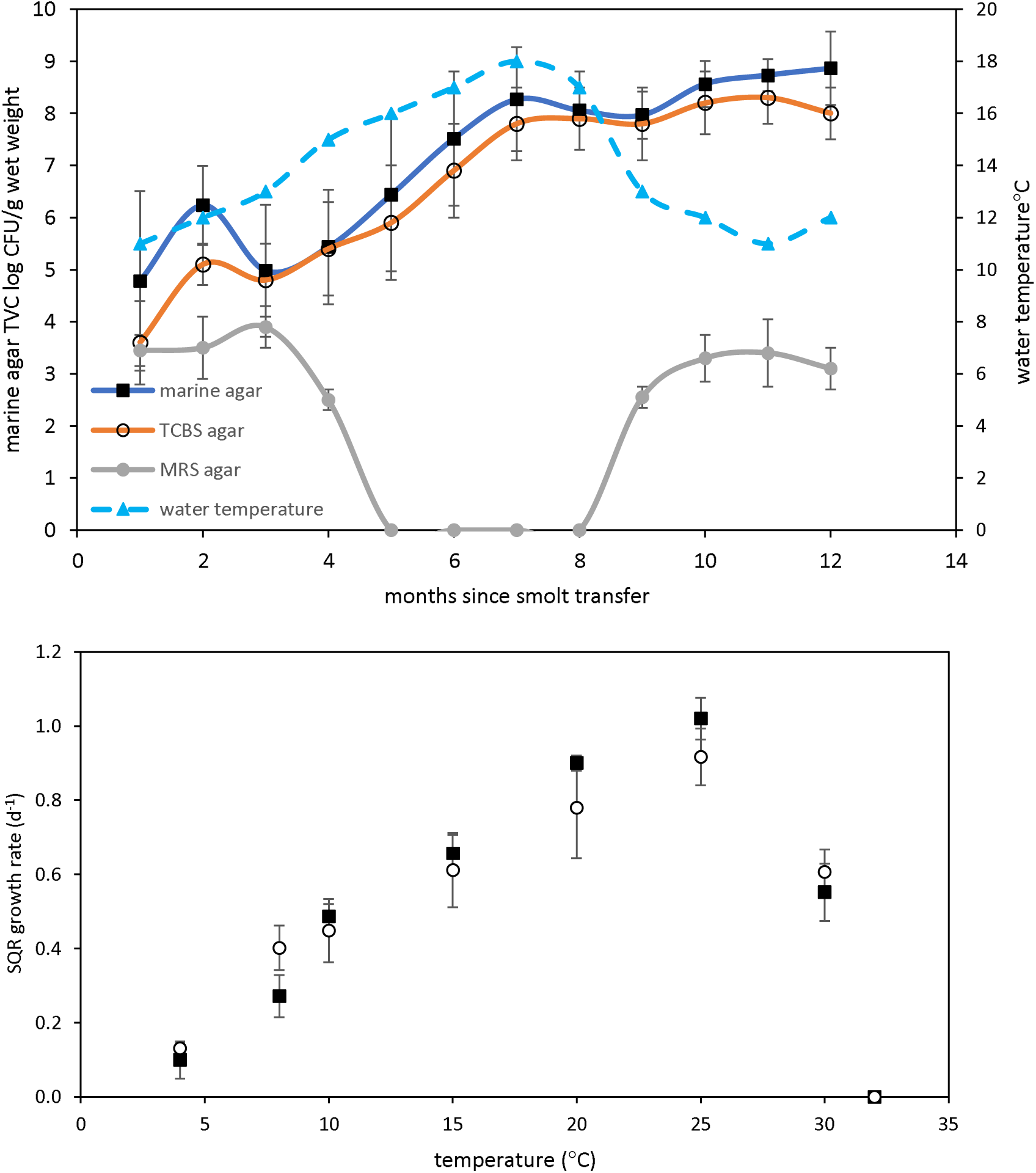
The top graph shows TVC fata for marine, TCBS and MRS agar (at 25 C) for bacterial populations obtained from Atlantic salmon digesta collected at different times after smolt transfer to marine farms. This is shown relative to water temperature. MRS counts were below detection limit during the summer months. The bottom graph shows the growth rates of salmon gut isolates at different temperatures. The black squares and white circles represent growth in marine broth and marine broth containing 1% (w/v) bile salts.

### Comparison with feed and water samples

On average 70% of reads were chloroplast and mitochondrial 16S rRNA from the plant components of the feed pellet ingredient mix. Vibrionaceae constituted 3% of bacterial reads in the feed pellets. The remaining 27% of reads were from other bacteria (Supplementary datafile 1). In water samples at farm-sites Vibrionaceae contributed 4.6% of total reads. *Aliivibrio* OTUs and *Vibrio scophthalmi* had low presence in water and feed samples (Figure 1, Supplementary datafile 1). Unclassified Mycoplasmoidaceae sequences were not detected in either the water or feed pellet samples.

### Cultivation-based data

Sequence based findings are supported by bacterial TVC data, measured from the smolt transfer period till the following winter for three of the survey years. TVC was determined on MA as a non-specific medium since the digesta content is saline (20-25 psu). These results were compared to TCBS agar counts since it is able to estimate more specifically family Vibrionaceae counts. The MA TVC data indicates that measurable populations reached a maximum level of 8 to 9 log colony forming units (CFU)/g sample wet weight within 7 months after smolt transfer (Figure 5). The close similarity observed between MA and TCBS count data (differing on average by 0.4 ± 0.4 log unit) indicates most of the TVC data come from bile salts-tolerant marine bacteria. Based on distinct colony morphologies isolates were obtained from the MA agar plates and purified. Growth on MRS agar was found to be negative during the summer months but reached 3-4 log CFU/g in the cooler seasons.

### Isolate data and proliferating OTUs

A total of 96 salmon isolates were collected and identified as *Aliivibrio* (68 isolates), *Photobacterium* (18 isolates), *Vibrio* (2 isolates), *Enterovibrio* (2 isolates), *Shewanella* (8 isolates), and *Psychrobacter* (1 isolate). The 16S rRNA gene sequence-based phylogenetic tree for these isolates is shown in the supplementary information (Figure S3). Comparisons were made between OTUs of the 8 surveys with the isolates. In the case of *Aliivibrio*. OTUs related to *A. finisterrensis*and two related unclassified species, represented the vast majority of *Aliivibrio* reads that showed evidence of proliferation. The predominant OTUs are shown in Figure 4. Genome data derived 16S rRNA gene sequences helped define species identities within the short sequence read data (Figure S4). Many other *Aliivibrio* OTUs were detected as shown in Figure 4, however these had comparatively lower abundance in all of the surveys. Most *Vibrio*-related OTUs that showed proliferation were related to the species *Vibrio scophthalmi. Photobacterium* OTUs that showed evidence of an increase in abundance over time included *Photobacterium piscicola*, *Photobacterium toruni* and OTUs that matched known species and or could not be classified at species level. Most reads of the unclassified Mycoplasmoidaceae group were represented by only a single OTU, which was distantly related to members of genus *Malacoplasma*.

**Figure 4.**
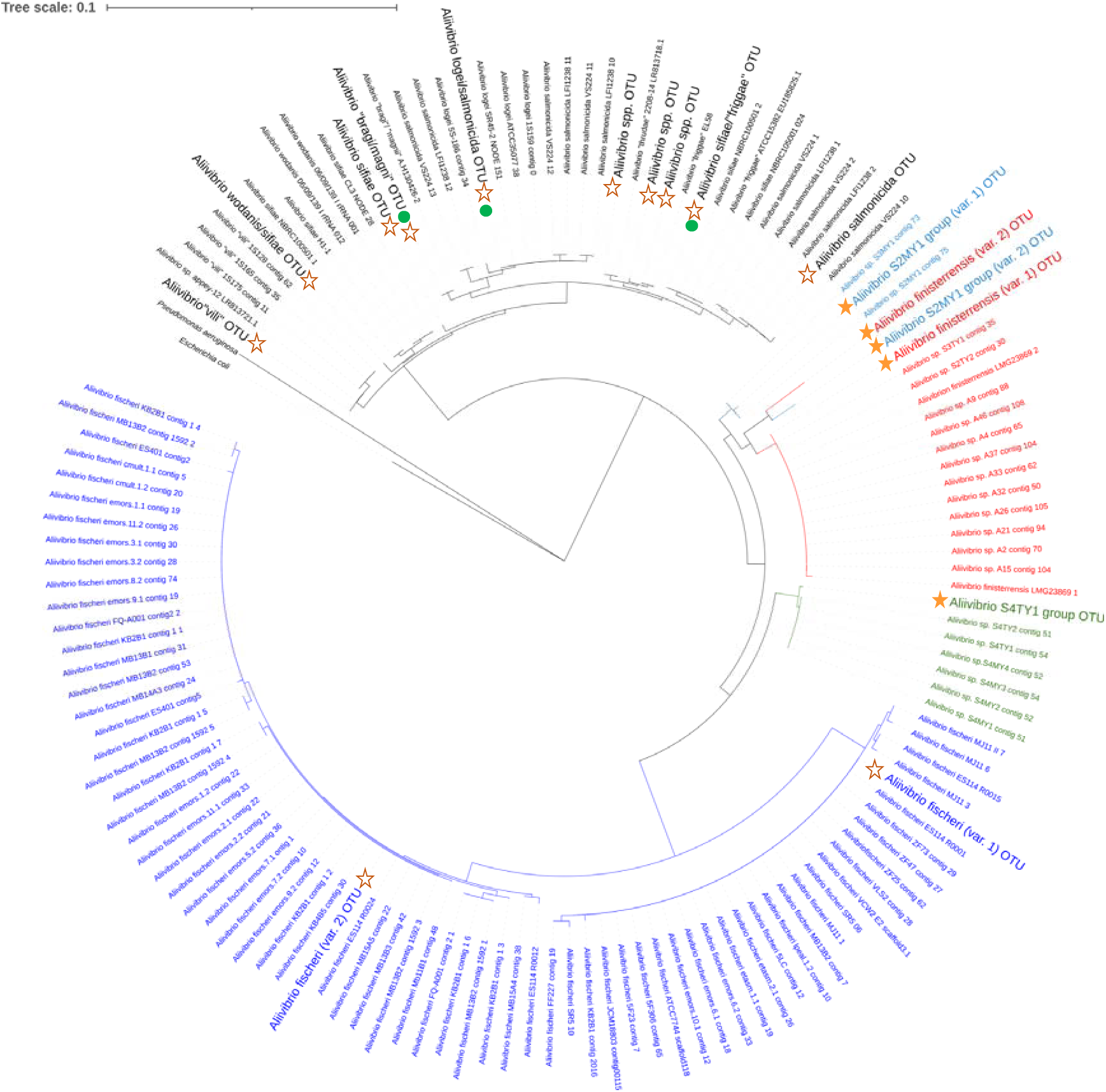
16S rRNA phylogenetic tree showing the relationship of *Aliivibrio* OTUs found across the 8 farm surveys. OTUs with filled stars were highly abundant in most surveys. OTUs with open stars were comparatively less abundant. Green circles donate the *Aliivibrio* species from the study of Vera-Ponce de Léon et al. (2024) based on alignment comparisons.

**Figure 5.**
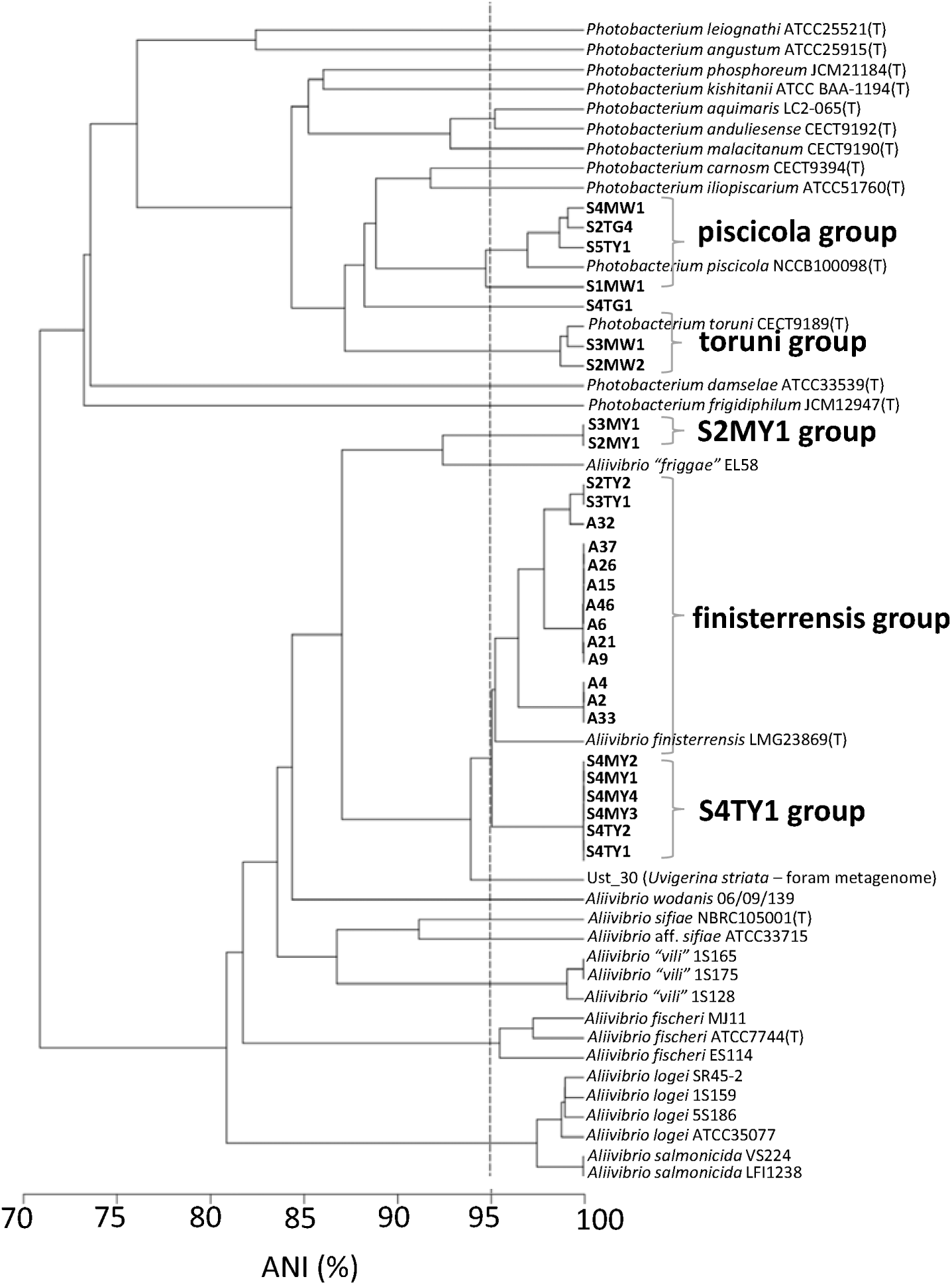
ANI-based unweighted pair mean group cladogram showing the taxonomic relationship of sequenced salmon isolates. The dashed line denotes the species cut-off of 95%. The range for species membership is considered to be in the 95-96% range.

### Genome sequencing of salmon isolates

Twenty-one of the isolates were genome sequenced. The *Aliivibrio* isolates that were sequenced comprised five ANI-defined genetic groups designated the A2, A9, A32, S2MY1 and S4TY1 strain groups (Figure 5). ANI and *in silico* DNA: DNA hybridization data (Figure S4) indicated the A2, A9 and A32 groups likely represent a single species (ANI >96% - Figure 6; >70% hybridization Figure S5). These three strain groups are most closely related to the type strain of *Aliivibrio finisterrensis* (LMG 23689^T^) based on ANI (95%) and in silico DNA: DNA hybridization (58-65%). The S4TY1 group is equidistant to the A2 group cluster and the *A. finisterrensis* type strain (94% ANI, 50-55% DNA: DNA hybridization), while the S2MY1 group is a more distinct *Aliivibrio* species with ANI at 80-87% with other available *Aliivibrio* genomes (Figure 5). The *Photobacterium* isolates belonged to either *Photobacterium piscicola*, *Photobacterium toruni* and an apparently novel species represented by strain S4TG1 (Figure 5). Strain S4TG1 had a 92% and 87% ANI to *P. piscicola* and *P. toruni* type strains, respectively. The other isolates obtained were most closely related to *Vibrio scophthalmi*, *Enterovibrio norvegicus*, *Shewanella pneumatophori*, and *Psychrobacter namhaiensis* (Figure S3).

**Figure 6.**
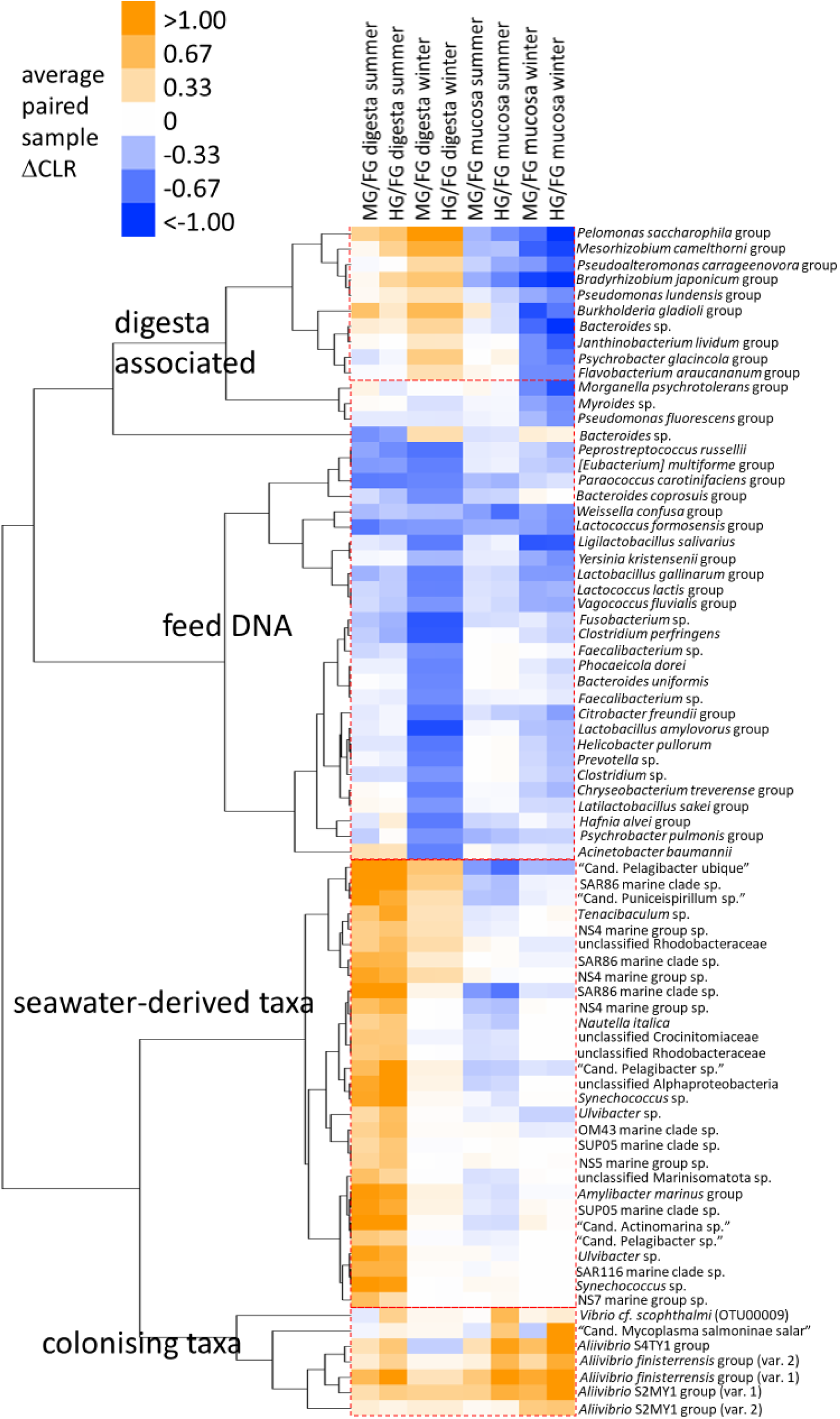
Heat map showing distribution of taxa comparing gut sections (FG, foregut; MG, midgut; HG, hindgut) for Atlantic salmon collected in the summer and winter of 2018. The sampling immediately followed feeding and was done over a 24 h period (Reid et al. 2024). The differences between abundances in these sections is indicated by the averaged differences in CLR values (ΔCLR) from paired samples (n=27 for each comparison). The OTUs included show significant values based on FDR adjusted significance testing and those showing large ΔCLR for at least one of the 8 dataset comparisons.

### Distribution of mucosa and digesta associated bacteria within the GI tract of farmed Atlantic salmon

Raw sequence data was utilised from the study of Reid et al., (2024) for an analysis of the distribution of bacteria through the extent of the digestive tract. For this experiment paired comparisons of abundances of OTUs was made between samples in the same fish. This was done between fore-gut versus mid-gut and fore-gut versus hind-gut digesta and washed mucosa. A total of 54 fish was examined this way with 27 fish sampled during summer and a second set of different fish sampled during winter. To do this comparison CLR values were compared for the same OTUs between paired samples resulting in the calculation of averaged CLR differences (ΔCLR). The OTUs included in the resultant heat map (Figure 6) had large ΔCLR values (>0.4 or <-0.3) and non-parametric effect sizes (*r* >0.5) as well as significant W-values (p<0.05, FDR) for at least one of the 8 datasets indicated. The OTUs included in the heat map in general represented the most abundant in the overall datasets being present in at least 20-25% of total samples. Less abundant OTUs generally were too inconsistent in distribution to obtain results that passed the criteria used for establishing significance. The results indicated a distinctive contrast between summer and winter samples based on hierarchical clustering patterns (Figure 6). In this respect four groups of OTUs was resolved. The first of these groups included OTUs termed “colonists” and included OTUs of *Aliivibrio finisterrensis* and other Aliivibrio OTUs belonging to the S2MY1 and S4TY1 groups. The predominant unclassified Mycoplasmoidaceae OTU and the most abundant *V. scophthalmi*-related OTU were also in this group. These species were defined as colonist species since they show positive ΔCLR values for mid- and hind-gut mucosa samples and are the same OTUs contributing to proliferation in the multiple survey digesta data (Figures 1 and 2). A second group of OTUs, which we refer to as “digesta-associated”, the most abundant being of the genus *Pelomonas*, demonstrate positive ΔCLR values for digesta samples but were practically absent in mucosa. A third group of consisting of taxa typical of seawater (e.g. SAR11, SAR86, *Synechococcus*, Rhodobacteraceae) show a similar response but increases in ΔCLR in the midgut and hindgut are restricted to the summer. The final “feed pellet DNA group” had reduced ΔCLR values for both digesta and mucosa samples for both summer and winter and mainly included OTUs grouped into Lactobacillales, Clostridia, Tissierellia, Bacteroidales, Enterobacteriaceae and Fusobacteriales.

### Salmon-associated *Aliivibrio* growth attributes

An analysis of the basic growth and phenotypic properties (Table 2) of the Atlantic salmon Aliivibrio strains was performed to assess environmental, growth and nutritional features. A focus was performed on *Aliivibrio* given these bacteria show year-on-year abundance increases and are associated with gut mucosa. The salmon isolates all have a mesophilic growth range (8 to 30°C) with an optimum at 20-25°C. In marine broth specific growth rates are (<0.6 d^-1^) below 10°C (Figure 3). All of the isolates are strictly halophilic. Optimal growth occurs with 2-3% NaCl or sea salts but no growth occurs with less than 0.5-1.0% NaCl. The strains are bile salt tolerant (1-3% ox bile salts) (Table 2). The strains were able to form biofilms on glass or polystyrene to different degrees, with only the S4TY1 and A2 group strains able to form significant biofilm on polystyrene within 2-3 d (Table 2). The isolates are non-bioluminescent except for two strains of the A32 group.

**Table 2.** Phenotypic data for *Aliivibrio* genetic sub-groups isolated from farmed Atlantic.

| Phenotype: | A2 group | A9 group | A32 group<sup>a</sup> | S4TY1 group | S2MY1 group |
| --- | --- | --- | --- | --- | --- |
| Growth temperature range (°C)<sup>a</sup> | 8-30 (25) | 8-30 (25) | <8-30 (25) | 4-30 (25) | 4-30 (25) |
| Salinity range for growth (sea salts % w/v)<sup>b</sup> | 1.0-5.0 (2.5) | 1.0-5.0 (3.0) | 1.0-5.0 (2.5-3.0) | 1.0-5.0 (2.5) | 1.0-5.0 (2.5) |
| pH range for growth<sup>c</sup> | 5.0-9.0 (6.5-8.3) | 5.0-9.0 (6.5-8.3) | 5.0-9.0 (6.5-8.6) | 5.0-9.0 (6.5-8.6) | 5.0-9.0 (6.5-8.6) |
| Maximum toleration of bile salts (% w/v)<sup>d</sup> | 2.5 | 2.0-2.5 | 3.0 | 1.0-2.0 | 2.0-2.5 |
| Bioluminescence | - | - | varies | - | - |
| Biofilm formation on polystyrene/glass<sup>e</sup> | ++/++ | -/++ | varies (- to ++)/++ | +++/+ to ++ | -/++ |
| Chitinolytic, carbohydrate fermentation | + | + | + | + | + |

<sup>a</sup>Detectable growth ( $A_{600} \geq 0.2$ ) occurring within 48 h incubation. The approximate optimal temperature for growth is shown in parentheses, doubling time ranged from 4.2 to 6.0 h at this temperature. Strains S2TY2 and S3TY1 showed growth at 4°C within 48 h but A32 did not.
<sup>b</sup>No growth occurs in the presence of $\leq 0.5\%$ or $\geq 7.0\%$ (w/v) sea salts; optimal salinity for growth is shown in parentheses.
<sup>c</sup>Values in parentheses denotes luxuriant growth occurs within the indicated pH range.
<sup>d</sup>Oxoid ox bile salts no. 3 LP0056 - 50% w/v cholate; 50% w/v deoxycholate. The value shown is the solution concentration that allows detectable growth ( $\geq 0.2 A_{600}$ nm).
<sup>e</sup>The degree of biofilm formation is based on the crystal violet assay absorbance ( $A_{550}$ nm) values for growth developed after 24 h of incubation at 25°C (+++ $\geq 0.6$ , ++ 0.3-0.6, + 0.1-0.3, - <0.1). Strain A32 did not form biofilms on polystyrene and is non-bioluminescent.

### Genome wide analysis for colonization-associated and toxin genes

The *Aliivibrio* salmon isolate genome data was annotated and investigated for genes that could be linked to colonization and virulence phenotypes. The analysis was aided by available research data on the function of genes in *Aliivibrio fischeri* (Chavez Dozal et al., 2012’ Aschtgen et al., 2019; Visick et al., 2021), *Aliivibrio salmonicida* (Skåne et al., 2022) and various *Vibrio* spp., in particular *V. cholerae* (Cho et al., 2021), all of which are good at forming biofilms in hosts. The results (Table 3) indicate the salmon isolates have a wide array of genes that can be linked to colonization, many of which are common across the genus *Aliivibrio* as well as the more distantly related species *Vibrio scophthalmi* (Table 3). In this respect all of the isolates possessed homologs for a range of adherence factors including lytic chitin monooxygenase (*gpbA*, LPMO10B), outer membrane colonization factors (*ompU*, *ilpA*, MAM7), chitin-regulated pili (*pilABCD*), and MSH-type pili (*msh* genes). Curli fibre genes (*csg* genes) and tight adherence pili (*tad* genes) are present in some of the strain groups. The isolates also have homologs for putative mucinases (*stcE*/*tagA*/*hapA* or *afcD*/*sslE* families). Most strain groups have syp-type, VPS-type and cellulose polysaccharide synthesis gene loci and genes associated with polysaccharide secretion (*eps* gene cluster). The isolates possess many genes known to be mechanistically required for biofilm formation (*lapV*/*lapI*, *lapBCDEDG*). Some of the isolate’s groups had homologs of biofilm associated genes that occur in *V. cholerae* but are absent in most other *Aliivibrio*. This includes rugosity factors *rbmABD* and *rbmC/bap1* that generate a thick, resilient biofilm. All of the isolates possess regulatory genes known to control polysaccharide and biofilm formation (*hapR*/*litR*, *vpsR*, *vpsT*, *hbtRC*). Furthermore, most strains also possess type VI secretion (T6SS) systems and usually have multiple associated effectors (e.g. *hcp*, *vgrG*). Interestingly, strain groups S2MY1 and S4TY1 have genes for the multi-subunit cytolethal distending toxin (*cdtABC*) (Table 3), which were most similar (50-55% amino acid identity) to genes found in pathogenic *E. coli* and *Salmonella enterica* (Figure S6). Known virulence determinants for Vibrionaceae were otherwise absent. Other *Aliivibrio* strains have a range of toxins that could be identified.

**Table 3.**
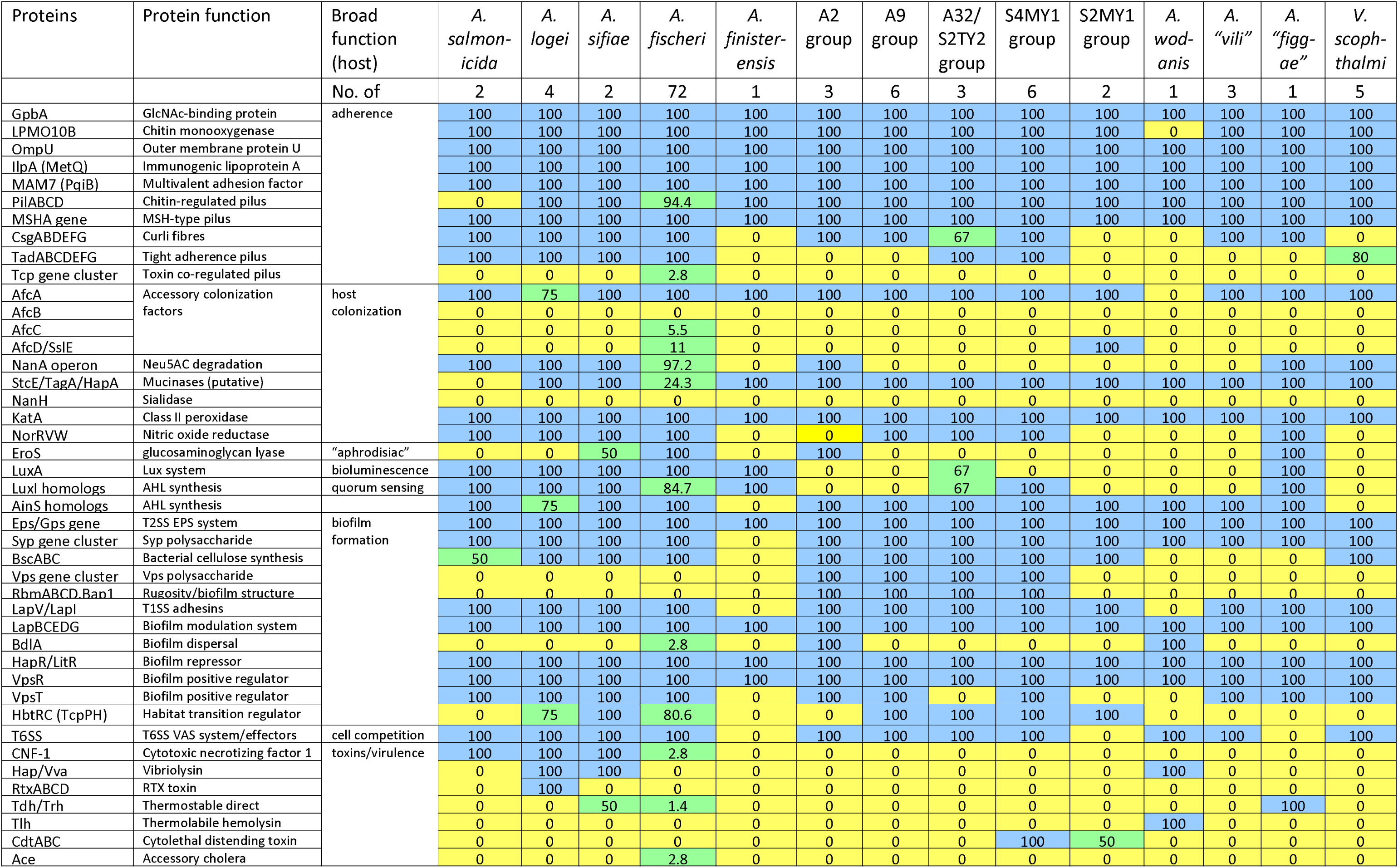
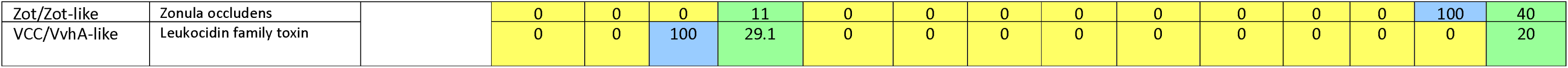
Host colonization relevant protein content in *Aliivibrio* species and *Vibrio scophthalmi* based on genome annotations and protein similarity analysis. Coloured values represent the percentage of strains containing the indicated protein homolog having >40% similarity and >80% of length to the reference protein. Footnote for Table 3. Of the 74 *A. fischeri* genomes 3 represent the type strain (ATCC 7744 = DSM 507 = JCM 18803) and are considered as one strain for the analysis. AfcB and AfcD homologs are present in most strains but only have 35-40% similarity with homologs from *V. cholerae*. and PilC homologs are absent in some *A. fischeri* and *A. salmonicida* strains. The *tad* operon may be present in two separate alleles, but three alleles is rare. MSHA genes *mshE*, *mshP* and/or *mshQ* may have diverged homologs in some strains. The *tcp* gene cluster occurs in the very similar *A. fischeri* ES114 and VCW2E2 genomes, but both lack *tcpR*. Some strains possess two or three StcE/Tag-like homologs. GAG (glycosaminoglycan) lyases of the PL8 family are present in *A. logei* IS159 and all three *“A. vili”* strains (1S128, 1S165, 1S175) though these proteins share only 38-40% homology with the GAG lyase EroS of *A. fischeri* ES114. The Vps cluster gene *vpsC* is not present in any strain while additional genes in the cluster are found in some strains e.g. strain A32. Many strains have a Zot domain-containing protein except ES114 and VCWE2E, which have adjacent Zot and Ace homologs.

## Discussion

Based on analysis of the main taxa present in Atlantic salmon GI tract digesta and mucosa world-wide the main taxa that predominate are analogous to what is observed in Tasmanian Atlantic salmon (Table S6). From this higher level view the presence of Vibrionaceae is most evident when surface seawater (or seawater tank) temperatures are above 12-13°C and include smolt or post-smolt being sampled weeks to months after deployment to a marine location (Godoy et al., 2015; Bozzi et al., 2021; Lorgen-Ritchie et al., 2021; Brealey et a. 2022; Schaal et al., 2023; Dhanasiri et al., 2023; Huyben et al., 2020; Heys et al., 2020, Kazlauskaite et al., 2021,2022, Li et al., 2021, 2022, Villasante et al., 2022). In the case of Tasmanian farmed salmon, we observed *Aliivibrio*, *Vibrio*, *Photobacterium* and unclassified Mycoplasmoidaceae were the only taxa that reliably increased in abundance over time over the 9-year survey period. Other taxa that showed increases were much less abundant and prevalent or often declined in abundance. The measured semi-quantitative increase of 2 to 3 log units (based on CLR data) and the MA/TCBS TVC data indicates this relates to a biomass increase in stripped fecal samples. The TVC data based on traditional agar provides credence for these findings by providing a population scale to work from. Based on this Vibrionaceae expand and predominate clearly by the late summer-early autumn a process that takes about 6 to 8 months. After which overall populations decline slightly but remain stable in the winter months following the summer/autumn peak.

The species of the main proliferating taxa, especially of genus *Aliivibrio*, were analyzed in detail. We obtained isolates of the predominant OTUs present for *Aliivibrio* as well as some *Vibrio* and *Photobacterium* and these were genome sequenced. These isolates were obtained from high dilutions of samples on MA and thus are highly abundant in the digesta samples (>7 log units/g for digesta from summer fish). The genome sequencing in revealing 16S rRNA gene variants aided the identification of these species in the MiSeq-based 16S rRNA gene community analysis. The study of Vera Ponce de Léon et al., (2024) identified a number of *Aliivibrio* that were predominant in a set of saltwater fish from a Norwegian coastal site, this included *Aliivibrio salmonicida*, *A. finisterrensis* and two novel *Aliivibrio* species. Based on alignment the novel species are most closely aligned to the placeholder “*A. bragi*”/”*A. magnii*” clade of Klementsen et al., (2021) while the other species is related to *A. sifiae*, *A. logei* and *A. salmonicida* group. Due to these bacteria hosting multiple 16S rRNA variants as well as the limitation of some species lacking sequenced type strains (namely *A. logei* and *A. salmonicida*) it is difficult to be more definitive in species classification. From the results here Tasmanian salmon contain quite different predominant *Aliivibrio* species. Most strains are related to A. finisterrensis- (A2, A9, A32 and S4TY1 groups) while a novel group (S2MY1 group) is most closely related to placeholder species *“A. friggae”* (Klemetsen et al., 2021). Collectively A2, A9 and A32 strains group were the the most abundant amongst fish while the other taxa are less abundant by 10-fold and 30-fold, respectively. Based on the ANI and in silico hybridization data these strains represent a single species that is closely related but distinct from *A. finisterrensis*if the species boundary guidelines is strictly interpreted, i.e. 70% DNA:DNA hybridization (Bach et al., 2023). However, the 16S rRNA genes of the strains are indistinguishable from the *A. finisterrensis* type strain. The main *Vibrio* species present in Tasmanian farmed salmon was also different to Norwegian salmon (Vera Ponce de Leon et al., 2024) with the species *V. scophthalmi* predominant instead of *V. splendidus*. Photobacterium found in Norwegian salmon seem to be mainly of the species *P. phosphoreum*, however in Tasmanian salmon the species *P. piscicola* and *P. toruni* were predominant instead, amongst a range of other potentially novel species. *Shewanella* and *Psychrobacter* seem to be relatively abundant as found in the de Leon and this study. The species-level differences could relate to many factors including sea surface temperature, the (epi)genetics of the salmon (Hansen et al., 2023) and an implicit amplification effect of strains becoming abundant over time the D’Entrecasteaux Channel region due to the high concentration of active Atlantic salmon farms. The heightened abundance means they are more likely to colonize than less abundant species as suggested previously (Heys et al., 2020).

We establish here that the proliferating taxa in the Atlantic salmon samples occur in abundance in washed mucosal samples and show an increasing abundance trend from the fore gut to the hind gut. The data analyzed included summer and winter fish cohort specimens since there was an interest in the investigating the effect of seasonal temperature differences on gut microbiome dynamics in more detail (Reid et al., 2024). In this respect *Aliivibrio* salmon bacterial isolates from Tasmanian salmon only grow relatively slowly below 10°C (<0.6 d^-1^, Figure. 5). This temperature is the minimum winter temperature recorded in the past decade in the main salmon growing region of Tasmania (Fig. S1). Based on the laboratory culture results the growth rates of *Aliivibrio* in spring and summer when water temperatures are >15°C should exceed 1.0 d^-1^. In winter water temperatures (∼10-13°C in southeast Tasmania) populations seem to be slightly lower than the summer peak possibly due to slower growth. Since warmer seawater conditions would enhance the growth rates of Vibrionaceae it would be expected more intense summer sea surface temperatures could promote the rapidity of the colonization process and result in greater biomass levels. It is unknown how the microbial diversity would be affected, for instance the appearance of only one predominant species, or instead there will be more stochasticity in the assembly of gut microbiomes in salmon. The colder waters typical for northern hemisphere salmon farming regions (i.e. Norwegian annual range is 11-17°C in summer and 4-5°C in winter), could explain the contrast in the species make up of Norwegian and Tasmanian farmed salmon. Based on our data, if water temperatures drops below 8-10°C *Aliivibrio finisterre*nsis and S2MY1 group related strains would be largely absent.

Examining the distribution of colonizing taxa in paired gut samples was useful in obtaining more targeted information on microbiome dynamics and structural relationships. Differences between mucosa versus the digesta were highlighted, as well as the effect of seasons. Usefully mucosal washed samples are not affected by the feed background as indicated by very low detection of chloroplast and mitochondria-derived 16S rRNA sequences originating from feed. On this basis the ΔCLR data indicates few taxa seem to actively colonize the mucosal layer in Atlantic salmon and include most but not all of the acknowledged proliferating taxa. By contrast digesta as would be expected possesses a very strong “diet signal” (Karlsen et al., 2022). The ΔCLR data from paired samples indicated the abundances of certain OTUs decline from fore-gut to the hind-gut. The decline in these OTUs when progressing the gut could represent dilution of feed-associated taxa. Members of order Lactobacillales (the lactic acid bacteria) were prominent amongst these declining OTUs. A heat map was created (Figure S7) comparing the overall most abundant OTUs in the feed, water and the digesta samples that came from the same hatchery and farm locations (the 2018 survey, table 1). Many abundant OTUs in digesta are the same as those from feed pellets. Some also occur at sporadic, low abundances in the water. The taxa included *Weissella confusa*t,he most abundant OUT in the feed pellets, *Streptococcus*, *Pseudomonas*, *Shewanella*, *Psychrobacter*, *Aeromonas*, *Myroides*, *Acinetobacter*, and an unclassified OTU belonging to family Bacteroidaceae. From this some OTUs could be just from feed pellets where the DNA or cellular remnants are abundant and possibly derived from bacteria added to feed ingredients as a probionts. Distinct colonies on MRS agar indicate that lactic acid bacteria are viable in digesta though populations are much lower than Vibrionaceae and could not be detected in summer samples. Despite this in the metabarcode sequence farm surveys Lactobacillales levels had a median proportion of 4% (range 0.02 to 53% of reads, Supplementary datafile 1). This dichotomy creates a problem in interpreting the actual nature of some taxonomic groups in 16S rRNA gene data derived from salmon digesta. Cultivation-based confirmation is required to get an accurate reflection of the gut microbiome or require advanced RNA-based techniques.

Another microbiome “artifact” that demonstrated interesting gut transect differences included seawater bacteria. A group of OTUs were found to show increased abundance in the digesta in the hind-gut relative to the fore gut digesta during summer but not winter. Distinctively the response between fore-gut and mid-gut was much more muted, did not occur for the mucosa samples, and did not occur in winter. Prominent amongst these bacteria were marine Alphaproteobacteria (SAR11, SAR116 clades), Gammaproteobacteria (SAR86 clade), Flavobacteriales (NS4, N5, NS7 clades), Actinomycetota (“Candidatus Actinomarina”) and Cyanobacteriota (*Synechococcus*). These taxa are typical seawater specialist taxa, which are strictly oligotrophic or in the case of *Synechococcus* obligately phototrophic, unculturable on agar media and have streamlined genomes. To date there is no evidence that such bacteria have direct functional relevance or impact in Atlantic salmon. A possible reason why seawater specialists are not only readily detected and show collective similar distribution patterns to each other is due to the high water content in the digesta samples. Digesta water content was found to be greater in summer fish than those examined during the winter (Reid et al., 2024). There is a distinct possibility that the abundance increase by seawater bacteria is due to water concentration effects. Since intestinal residence time is longer during winter (Mock et al., 2022) there is more time for water absorption within the gut especially in the foregut (pyloric caeca). The “drinking” rate is increased in salmon adapted to seawater and during active feeding and is used for the purposes of osmoregulation (Eddy 2007) maintaining a level of salinity in the gut lumen. The effect of water uptake could lead to seawater bacteria having a greater overall representation in the microbiome data. This occurrence has also been suggested as a reason that seawater dwelling Vibrionaceae are also able to colonize the gut (Bjorgen et al., 2020) since the salinity is suitable for their growth with most species being halophilic (Table 2). The indicated seawater specialists are common in Tasmanian waters, especially during summer (Brown et al., 2024). Many of the seawater taxa become less abundant in winter in Tasmanian waters. These are replaced by the growth of ammonia-oxidizing archaea and rhodopsin-containing heterotrophic archaea, which are not detectable with the primers used in the study.

Since most sea-going wild salmon and those that have returned to rivers also possess considerable levels of *Vibrionaceae* (Supplementary data Table S1, Llewellyn et al., 2016), colonization by Vibrionaceae is clearly a natural process. The observation that *Aliivibrio*, unclassified *Mycoplasmoidaceae* and *Vibrio scophthalmi*-related strains are able to colonize the mucosa suggests these taxa possess traits compatible with salmon colonization. The colonization process may involve a chemotactic element and is not simply a random process of water transfer, however the details remains to be determined.

We have hypothesised previously *Aliivibrio* colonizers are a possible cause of the production of fecal casts in fish sampled during summer (Reid et al., 2024). Fecal cast production in finfish has been attributed to disease or stress (Breen et al., 2021, Dhar et al., 2017, Birrell et al., 2003, Figueroa et al., 2017, Skår et al., 2007). The expected higher growth rates and biomass of *Aliivibrio* and *Vibrio* spp. in summer may increase the risk of dysbiosis or disease, but this relies on the species having some capacity to affect the host negatively. At this point in time the causal relationships remain unresolved. As a preliminary attempt to understand these responses we surveyed the genomes of the salmon isolates to find possible colonization- and virulence relevant mechanisms. Since *Aliivibrio* spp. predominate and demonstrate a capacity for colonization of the mucosal layer we particularly focused on this genus for understanding colonization mechanisms. *Vibrio scophthalmi* (and/or closely related species) genomes were also compared utilizing data available from the NCBI database. Based on the annotation results most if not all *Aliivibrio* possess an array of traits that are likely the basis for adherence and subsequent colonization in fish and other marine animal hosts.

A near universal trait for Vibrionaceae is motility, which is needed for initial adherence in the gut as well as spreading throughout (Homma et al., 2022). The salmon associated bacterial strains examined were all motile and based on the genome data this is only driven by polar flagella.

Homologs of flagellin genes forming lateral flagella (*laf* genes found in *Vibrio vulnificus*), which could aid spread in viscous fluids, in or on the host (Homma et al., 2022) were not found in any *Aliivibrio* genome. Vibrionaceae also produce several types pili and a range of other specific proteins to enable stable adherence to host gut cells and mucosa. Lytic chitin monooxygenase proteins LPM010A and LPM010B (Skåne et al., 2022, Wong et al., 2022], chitin-regulated pili (Paranjpye and Strom 2005), an MSHA-type pili (Teschler et al., 2015), were found in all *Aliivibrio* strains and have been shown in *A. fischeri* to be mechanistic elements for adherence and establishing populations in fish and squid host GI tracts (Visick et al., 2021). Most of the salmon isolates also possessed genes for curli fibre synthesis (*csg* operon) (Karan et al., 2021) while 1 or 2 alleles for the tight adherence pili (Tad operon) (Pu & Rowe-Magnus, 2018) were present in specific strain groups (A32, S4TY1 groups). Toxin co-regulated pilus genes (*tcp* operon) (Oki et al., 2022) were only found in a minority of *A. fischeri* strains. Potentially other proteins could be involved as well for adherence as found for *V. cholerae* and that are broadly conserved in Vibrionaceae e.g., IlpA/MetQ, OmpU, MAM7/PqiB (Lee et al., 2011; Goo et al., 2006; Yang et al., 2018; Krachler et l. 2011) (Table 3). A homolog of outer membrane accessory colonization factor gene *afcA,* which influences spreading in the host (Hughes et al., 1995; Cai et al., 2018), is universally present. Homologs of the other accessory colonization genes (*afcB*, *afcC*, *afcD*), putatively linked to mucin chemotaxis and degradation(Valiente et al., 2018) are not evident in most *Aliivibrio* strains, however, diverged homologous genes (35-40% identity) do occur, for example homologs of *afcD*. Most of the salmon isolates possess a number of putative mucinases, including YghJ/SslE/AfcD (Nesta et al., 2014, Szabady et al., 2011) and/or StcE/TagA (Rossiter et al., 2017) family lipoprotein metallopeptidases. Adding to a probable capability of colonizing and degrading mucin and other proteins making up the gut epithelial layer are putative chondroitinase -like enzymes (polysaccharide lyase family 8). These glycosaminoglycan degradative enzymes are co-located with genes coding enzymes for D-glucuronate, N-acetyl-D-glucosamine and N-acetyl-D-galactosamine degradation in several *Aliivibrio* species. Chondroitinase activity (*eroS* gene) by *A. fischeri*has been shown to induce sexual reproduction in a choanaflagellate (Rossiter et al., 2017; Woznica et al., 2017), but the any role this capability has in gut colonization remains to be shown. The *eroS* gene and associated metabolic pathway genes are strain group specific being absent in most of the salmon isolates, except the A2 group. The potential ability to utilize N-acetylneuraminate (*<u>nanA</u>* operon) (Boyd et al., 2015) and other carbohydrates liberated from mucins (N-acetyl-D-glucosamine, D-galactose, L-fucose) is also present in some *Aliivibrio* species. Amongst the salmon isolates the *nanA* operon is found in the A2 group strains. All genomes surveyed have a *katA* homolog shown to be needed by *A. fischeri* for dealing with host-driven peroxide defenses (Visick & Ruby 1998) and also possess nitric oxide reductase (NorRVW), which could act to protect cells from host NO synthesis (Gardette et al., 2020)

Biofilm formation and compatibility with the host is likely key for colonizing bacteria to establish stable populations in fish. Bacteria not native to the host, such as commercial probiotic strains, may only colonize transiently and much less efficiently due to inability to form a stable biofilm (Li et al., 2018). As mentioned above model species, especially *A. fischeri*, were used here due to research on its ability to form biofilms in marine fauna (Fung et al., 2024). From this knowledge the salmon strains have genes that code proteins required for biofilm establishment, build-up and dispersal as found in *A. fischeri* (Christensen et al., 2020). Fundamentally the mechanisms enable populations to expand *in vivo* since the colonizing bacteria needs to spread spatially on mucosal surfaces. As found in *A. fischeri* and *Aliivibrio* in general, the salmon isolates have a type I secretion system (T1SS) which delivers the adhesin LapV and proteins of the Lap adhesin delivery system (operon *lapBCEDG*) (Christensen et al., 2020). In addition, the salmon isolates also possess a conserved T2SS for exopolysaccharide (EPS) delivery. EPS in the salmon isolates like that of *A. fischeri* includes the syp polysaccharide, coded by the *syp* gene cluster which includes defined regulators (e.g. *vspR*) (Dial et al., 2021). Similarly, conserved regulators are present for biofilm matrix formation, such as the quorum sensing/competence repressor HapR/LitR (Bjelland et al., 2012; Cohen et al., 2021). These genes are prevalent in most *Aliivibrio* species and homologs also occur in *V. scophthalmi* suggesting a degree of conservation of the fundamental biofilm formation mechanism. The gene locus of the syp polysaccharide was found to be syntenic (in relation to gene loci in *A. fischeri* strains) in the salmon strains as well as most other *Aliivibrio* species.

Beyond conserved traits, several genes were noted to be specific to species or sub-species strain groups (Table 3). These traits likely contribute to colonization in several ways. These could determine host specificity, an area that remains largely unresearched. For example, cellulose synthesis genes (*bsc* operon, Abidi et al., 2022) are present in genomes of 75% of *Aliivibrio* genetic groups while homologs for the VPS polysaccharide locus of *V. cholerae* (Schwechheimer et al., 2020) were found only in the genomes of salmon-colonizing *Aliivibrio* isolates. The VPS polysaccharide contributes to a viscous polysaccharide that enables *V. cholerae* and other *Vibrio* spp. colonization of a range of ecosystems and hosts (Fung et al., 2024). Some genes in this locus are different in the salmon isolates compared to *V. cholerae* and the synteny is not fully preserved, suggesting the polysaccharide formed conceivably could have different chemical properties. Furthermore, the same strains possess adjacent genes that may regulate biofilm thickness and structure as found for *V. cholerae* (*rbmABD*, *rbmC/bap1*) (Fong & Yildiz 2007). Many of the indicated genes are regulated by quorum sensing systems controlled by synthesis of different acylated homoserine lactones (AHLs) and autoinducer-2 (LuxS system) (Lupp & Ruby 2004). The salmon strains all possess *luxS* and possess genes for two AinS, LasI or LuxI family AHL synthetases (Table 3). This suggests the salmon isolates (and most other *Aliivibrio*) may be able to form an array of quorum sensing-related chemical messengers as found for *A. fischeri*(Girard et al., 2019).

Virulence gene distribution was of considerable interest, in particular to discover genes that might be linked to fecal cast production. The survey conducted examined genes flagged generically as virulence factors but also cross-referenced this with Vibrionaceae-associated proteins known to directly cause cytotoxicity of some form. This includes T6SS systems that have been suggested to sometimes be involved in host virulence as well as interbacterial competition (Rubio et al., 2019). In this respect, all of the salmon strains possess a T6SS system, except the S2MY1 group, and also possess multiple associated effectors (i.e., *hcp*, *vgrG*) (Crisan & Hammer, 2023; Cohen et al., 2023). Strains A9 and A21 possess both *hcp* and S-type pyocin-like (Scholl, 2017) homologs on a contig that is likely a large plasmid. T6SS is prevalent amongst most other *Aliivibrio* species. It is so far unknown if T6SS has any effect on hosts directly, however the T6SS in *A. fischeri* is used to effectively to become predominant in squid hosts (Guckes & Miyashiro 2023). By comparison toxin genes are sporadically distributed across *Aliivibrio*. For example, a minority of *A. fischeri* strains possess genes for CNF-1, Tdh, Ace, VCC/VvhA-like leucocidin and/or Zot/Zot-like proteins found in various pathogenic *Vibrio* (Cai et al., 2018). Surprisingly, *cdtABC* genes coding the subunits for CTD (Lai et al., 2021) occur in one S2MY1 group strain (S2MY1, but not S3MY1) and all six of the S4TY1 group strains. The gene nucleotide *and* the amino acid sequences for the three Cdt subunits were surprisingly found to be identical between the isolates and are distinct from homologs of *E. coli* and *Salmonella* (∼50-55% amino acid homology) and are also distinct from homologs found to occur in a single *V. cholerae* isolate 633012 (supplementary data Figure S7). The genes occur in a region of chromosome II where extensive transposase/integrase gene decay has occurred. The most populous salmon strain groups (A2, A9 and A32) completely lack any known genes linked to a known cytotoxic function.

The results indicated *Aliivibrio* possess various colonization-relevant genes that are group and species-specific. This suggests that colonizer genetic groups could have different colonization specificity and preferences. Many features are also shared with *Vibrio scophthalmi*, a very successful colonizer of fish that is sometimes an opportunistic pathogen (Zhang et al., 2020). The differences in this gene distribution may affect adherence, mucin/mucosal interactions, signal transduction, and biofilm structure within different hosts thus determining specificity, success rate and persistence of colonization. At this stage the genome data only offers a surface view, the active mechanisms of colonization are presently unresolved. The data provided in this study suggests a series of proteins and phenotypes that could be further researched in this respect. For example, putative mucinases were detected in the salmon colonizers. The *Vibrio cholerae* HapA mucinase facilitates penetration and detachment into the mucosal layer and also may release substrate sugars and amino acids (Silva et al., 2006). Few of the Atlantic salmon isolates seem to be able to degrade neuraminate, a major glycocomponent of intestinal mucins of Atlantic salmon (Jin et al., 2015). Mucinase activity by colonizer strains could instead mainly allow persistence in the mucosal layer (Silva et al., 2006). However, the colonizer community combined mucinase secretion activity could liberate a range of glycopeptides, amino acids, and sugar residues that can be used as nutrients by the microbiome as a whole. In this respect all colonizer strains could grow on chitin and possessed chitinolytic enzymes (e.g., LPMO10A, LPMO10B, Table 3) shown to be important in mucosal layer adherence and required for virulence in some strains (Skåne et al., 2022, Wong et al., 2012). Gut mucins of Atlantic salmon typically contain N-acetyl-D-glucosamine (Berktander et al., 2019) and it seems some chitinase-like enzymes are able to remove these residues from mucin (Skåne et al., 2022). Salmon mucin is also rich in galactose residues and from the genome data most of the salmon isolates were found to possess the Leloir galactose metabolism pathway based on the genome annotations.

Interestingly, salmon isolate groups also synthesize multiple types of polysaccharides that may be involved in colonization of marine animals and abiotic surfaces (syp, VPS and cellulose, Table 3). It is possible these polysaccharides co-contribute during colonization of fish. *Aliivibrio* genetic diversity may also impact other biological aspects of colonization, such as quorum sensing regulation. The types of AHL synthetases differed between the colonizer genetic groups and in some cases evened differ within the same group, for example the A32 group strains have either LuxI or AinS family AHL synthetases. In terms of the gut ecosystem *Aliivibrio* strains may be able to respond to the signals from other strains allowing rapid responses in gene regulation simultaneously maximizing fitness and competitiveness during both colonization. Potentially linked to signal transduction are pathogen-like functions by which some colonists exploit host cells. Virulence gene occurrence amongst *Aliivibrio* species, however, is quite sporadic and less conserved than other traits examined (Table 3). The occurrence of CDT-gene positive strains in some salmon isolates is intriguing but raises many questions. Are the genes expressed *in vivo*? Do they have a functional role that does not involve harming the host – for example aiding persistence in the salmon gut or are they merely unstable genes derived from recent horizontal gene transfer events? The distribution of CDT is presently unknown in marine bacteria but is likely extremely rare. The fact that the genes are identical between two different *Aliivibrio* genetic groups suggests lateral gene transfer involved the same source and occurred in a similar time frame. The genomic region, the *cdtABC* operon was found to be rich with pseudogenes, mainly remnant transposase genes. This suggests the transfer of genes could have been within a mobile element that is now decayed. Further research is clearly needed that involves linking fish and bacterial biology in controlled experiments to further understand how *Aliivibrio* and other colonizing taxa interact with Atlantic salmon. This study suggests several possible pathways for such ventures including adherence and biofilm formation, AHL-based signal transduction, mucosal interactions (mucinase production), competition between gut microbiota and possible impact on host cells by T6SS effectors, and finally potential toxin production by specific strains (e.g. CDT toxin).

### Conclusions

We found specific Vibrionaceae occur in high abundance in the GI tracts of Atlantic salmon farmed in Tasmania. Tough the same genera are present we found the species differed to what is generally seen in northern hemisphere farmed salmon. The gut profile survey revealed a restricted range of species, mostly novel, are robust colonizers of the mucosal layer. Such strains along could be a focus for future studies related to Atlantic salmon gut function and health especially pertaining to climate change. This owes to the colonization patterns of Aliivibrio seeming to be largely driven by water temperature. The most common *Aliivibrio* colonists lacked any obvious virulence factors, which is good news but at the same time other relatively abundant species were found to have specific virulence genes, including remarkably genes for the CDT toxin. Understanding the significance of such genes in relation to Atlantic salmon health and productivity as well as the biology of gut colonization is a logical next step. Combining temporal sampling with the genome biology of colonizing isolates proved very useful in defining the dynamics of colonizing bacteria and also defined traits that may be relevant to salmon health and welfare. The findings revealed interesting and complex traits amongst different *Aliivibrio* spp. genetic groups that likely contribute to their success in colonization of salmon.

## Supporting information

Supplementary datafile 1

Supplementary datafile 2

## Abbreviations

CDT: cytolethal distending toxin
RAS: recirculation aquaculture systems
OTU(s): operational taxonomic unit(s)
CLR: centered log ratio
CFU: colony forming units
ANI: average nucleotide identity
GGDC: genome-to-genome distance calculator
ΔCLR: centred log ratio difference (between paired samples)
LPMO: lytic polysaccharide monooxygenase
VPS: Vibrio polysaccharide (of *V. cholerae*)
MSHA: mannose-sensitive hemagglutinin (pili/fimbriae)
T1SS, T2SS, T6SS: type (I, II or VI) secretion system
AHL: acylated homoserine lactone
BLAST: Basic Local Alignment Search Tool
MA: marine agar
TCBS: thiosulfate citrate bile sucrose agar
PANNZER: Protein Annotation with z-Score
KEGG: Kyoto Encyclopedia of Genes and Genomes
VFDB: Virulence Factor Database

## Availability of data and materials

The raw 16S rRNA gene sequence data and metadata files are deposited at the NCBI SRA database under the BioProject codes listed in Table 1. *Aliivibrio* and *Photobacterium* genome sequence data are deposited in the NCBI database under WGS project codes WBVP00000000 (*A. finsterrensis* LMG 23689), SEZJ00000000 to SEZQ00000000 (*Aliivibrio* strains A32,A46, A37, A26, A21, A15, A9, A33, A4, A2), JARACN00000000 to JARACX00000000 (*Aliivibrio* strains S2MY1, S3MY1, S4MY1, S4TY1, A6, S2TY2, S3TY1, S4MY2, S4MY3, S4MY4, S4TY2), and JAYXUB00000000 to JAYXUK00000000 (*Photobacterium piscicola* strains S1MW1, S2TG4, S4MY1, S5TY1; *P. toruni* strains S2MW2, S3MW1; *Photobacterium* sp. S4TG1).

## Declarations

## Acknowledgements

The authors would like to thank Tassal Group Pty Ltd and Huon Aquaculture Pty Ltd for access to farms for sample collection. The authors would like to thank staff at Taroona EAF (Institute of Marine and Antarctic Studies, University of Tasmania) for tank set up and monitoring.

## Ethics approvals

Animal experimentation performed in this study and data obtained from previous studies has been approved under appropriate permits issued by either the University of Tasmania or the Tasmanian Government. Permit codes include DPIPWE 30/2009-10, A0012001, A0013971, A0015452, A0016307, A0016588, and A0017010.

## Supplementary figures and tables

**Figure S1.** Location of farms from which Atlantic salmon were sampled.

**Figure S2.** Proportion of taxonomic groups across the survey data including water, feed, and gut samples.

**Figure S3.** Identification of Atlantic salmon bacterial gut isolates.

**Figure S4.** Aligned 16S rRNA variants within the V1-V3 region for available *Aliivibrio* species including those from this study.

**Figure S5** In silico DNA:DNA hybridization between *Aliivibrio* salmon isolates and other *Aliivibrio* strains via the GGDC model 2 method

**Figure S6.** Comparison of concatenated cytolethal toxin subunit CdtA, CdtB and CdtC protein sequences from Atlantic salmon *Aliivibrio* isolates with Gram-negative pathogens possessing Cdt.

**Supplementary datafile S1.** Read counts for genera and unclassified gut microbiome taxa across multiple farmed Atlantic salmon surveys, feed pellets, hatchery and farm-site water in Tasmania, Australia.

**Supplementary datafile S2.** Differential CLR values across time for gut microbiome taxa across the surveyed duration at marine farms in Tasmania, Australia.

