## Supplementary material for "Characterization of bacteria colonizing the mucosal layer of the gastrointestinal tract of Atlantic salmon farmed in a warm water region": Figure S1

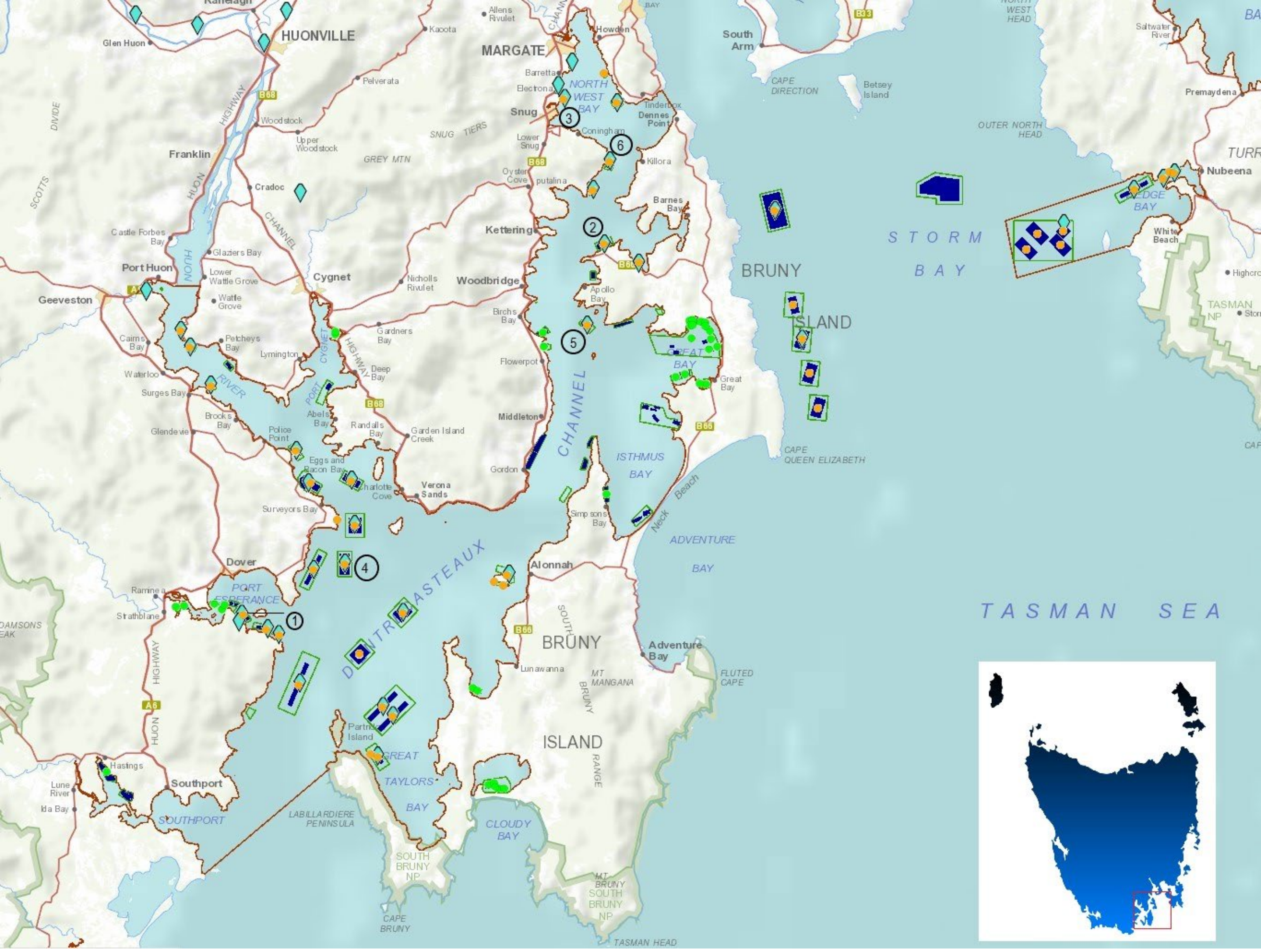


**Figure S1.** Location of farms from which samples were collected for this study and antecedent published studies as shown in Table 1. Farms sampled were at (1) Meads Creek, Port Esperance; (2) Robert's Point lease, D'Entrecasteaux Channel, (3) North West Bay, ); (4) Red Cliffs, D'Entrecasteaux Channel; (5) Soldier's Point, D'Entrecasteaux Channel; Sheppard's lease, D'Entrecasteaux Channel (6)

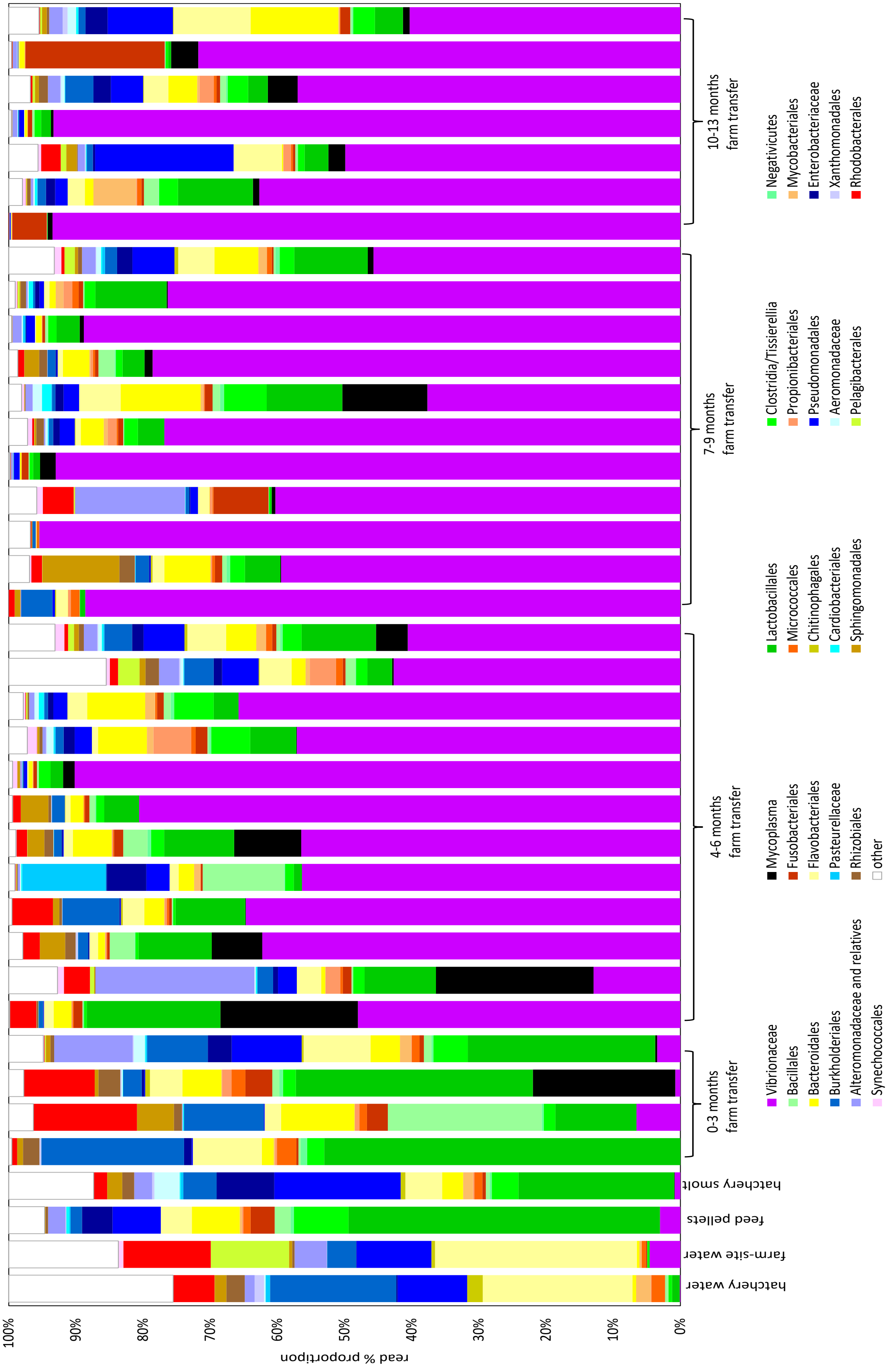


**Figure S2.** The proportional representation of taxa from farmed Atlantic salmon fish gut samples collected over several surveys in Tasmania (Table 1). More details are also shown in Supplementary datafile 1.

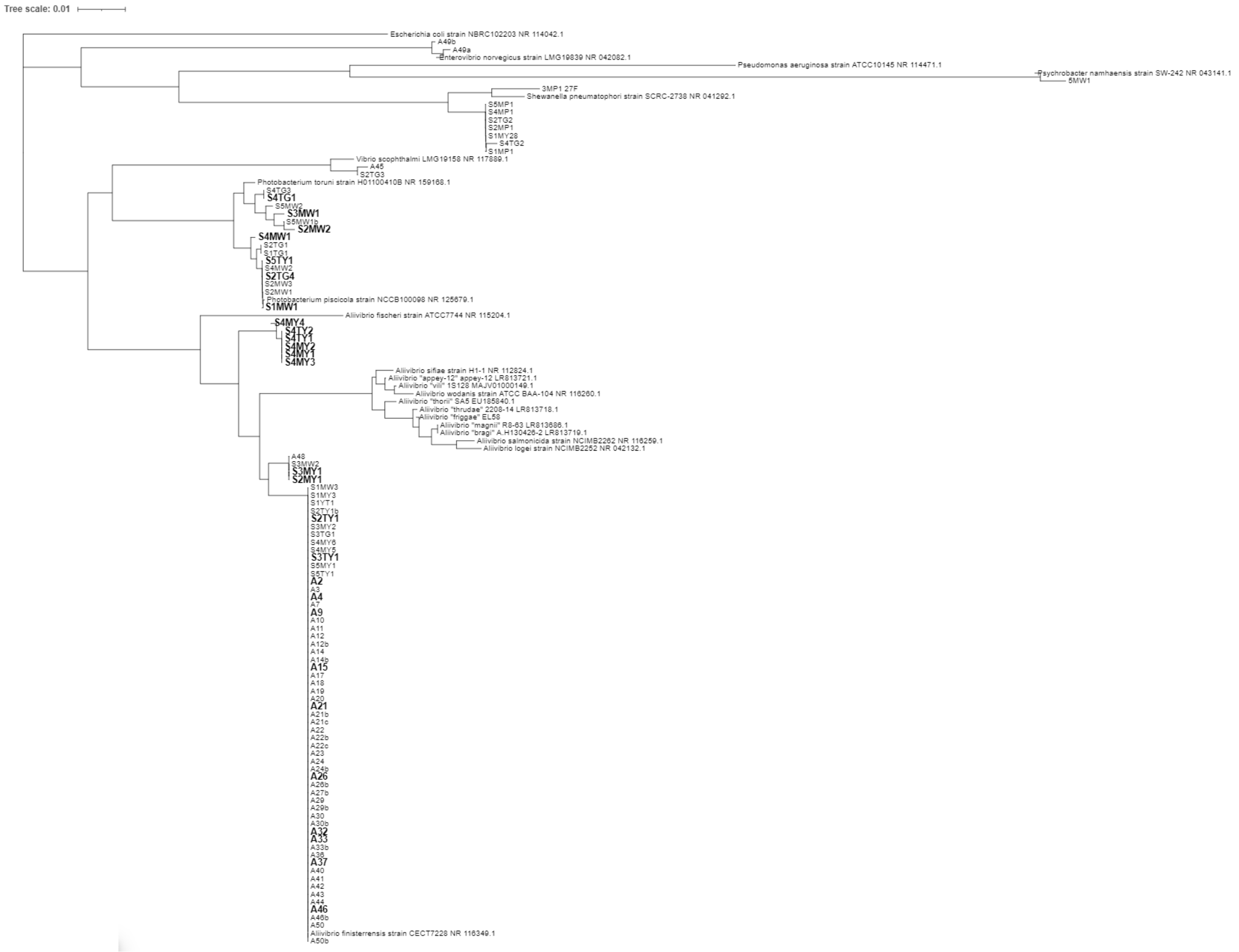


**Figure S3.** 16S rRNA based phylogenetic tree showing the closest related validly described species to bacterial strains isolated from farmed Atlantic salmon from south east Tasmania. Strain designations in bold were subsequently selected for genome sequencing.

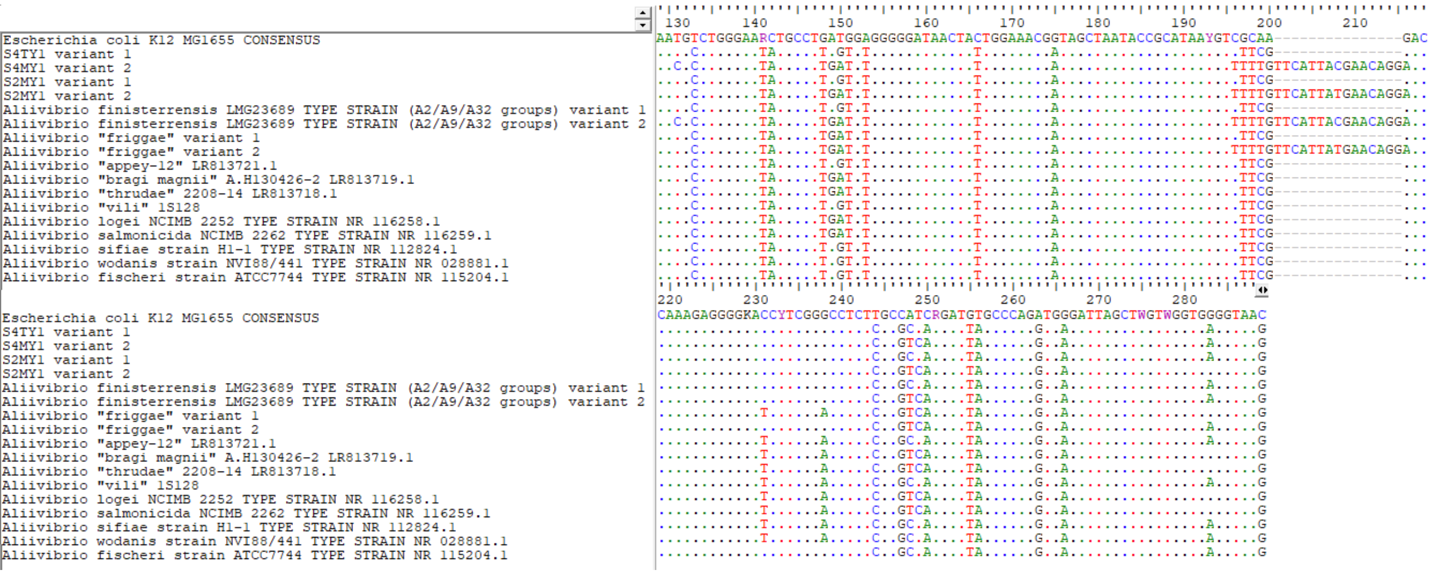


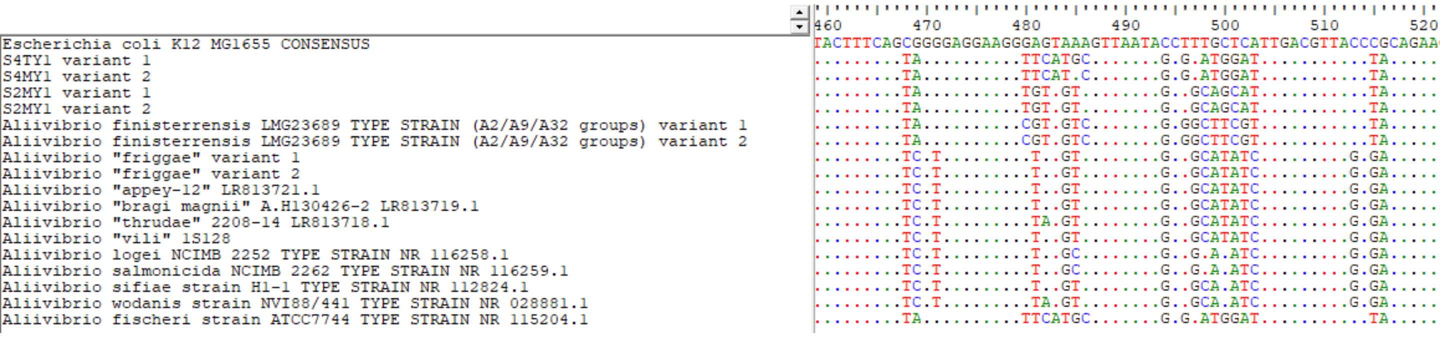


**Figure S4.** Alignment of sections of the V1-V3 region of 16S rRNA genes (*E. coli* numbering equivalent 10 to 534) for Aliivibrio species. The two main variants are shown for 3 major Atlantic salmon gut isolate groups (A. finisterrensis, S2MY1, S4TY1) as well as showing regions of hypervariability in the V1-V3. This resulted in OTUs

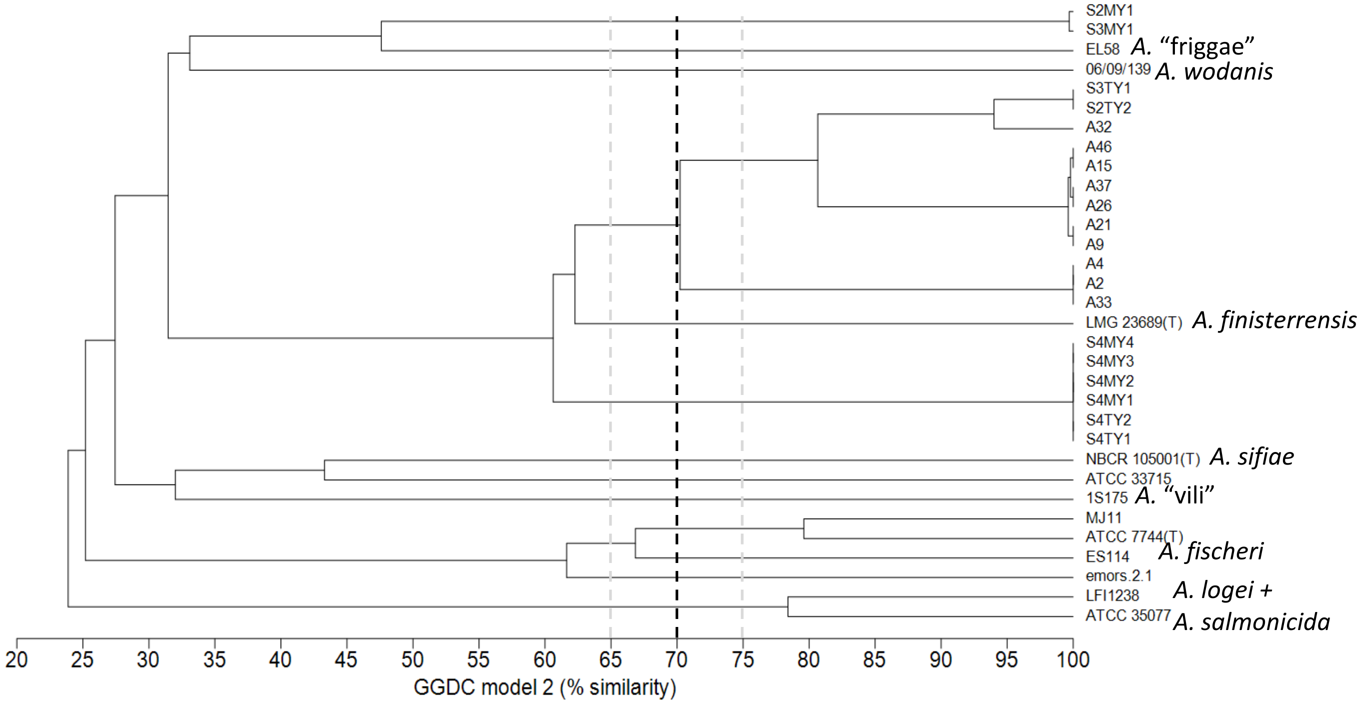


**Figure S5.** Group average-based tree based on similarities between *Aliivibrio* strain genomes. The similarities were generated with Meier-Kolthoff et al. GGDC model 2, which simulates *in silico* DNA:DNA hybridization. The classic demarcation for species boundaries of 70% is shown as a dashed line. The main features of the plot is that the A2, A9, and A32 groups cluster at the 70% boundary while the type strain of *A. finisterrensis* (LMG 23689$^{T}$) clusters with these groups at 58 to 65% similarity. The S4MTY1 group clusters with LMG 23689$^{T}$ at only 54% and 60-62% with the A2/A9/A32 groups. Strains of *A. logei* and *A. salmonicida* (neither are the actual type strains) cluster at 78%, while *A. fischeri* strains cluster at 60 to 80%. Group S2MY1 clusters closest to A. “friggae” EL581 at 43% similarity. The 60% similarity level approximately represents 94-95% average nucleotide identity. As a result, conservatively, the A2/A9/A32, S4TY1 and more obviously S2MY1 strains groups all represent separate species, however *A. fischeri* may not be a single species if this stance is taken. This situation is suggested in the GTDB database (gtdb.ecogenomics.org) where 6 strains (emors.2.1 – the placeholder representative, emors.5.2, emors.5.1o, emors.2.2, emors.5.1t, emors.1.2) are designated as *A. fischeri*_A while the 71 other strains form *A. fischeri sensu stricto*. Strain emors.2.1 (as shown above) has 60% similarity with the *A. fischeri* type strain ATCC 7744$^{T}$. GTDB uses an ANI circumscription radius of 95% to define a species though to retain as many species names as possible this boundary ANI values up to 97% are used. Increasingly ANI is used to demarcate bacterial species but this needs to be related to other data due to the fuzzy nature of bacterial species boundaries.

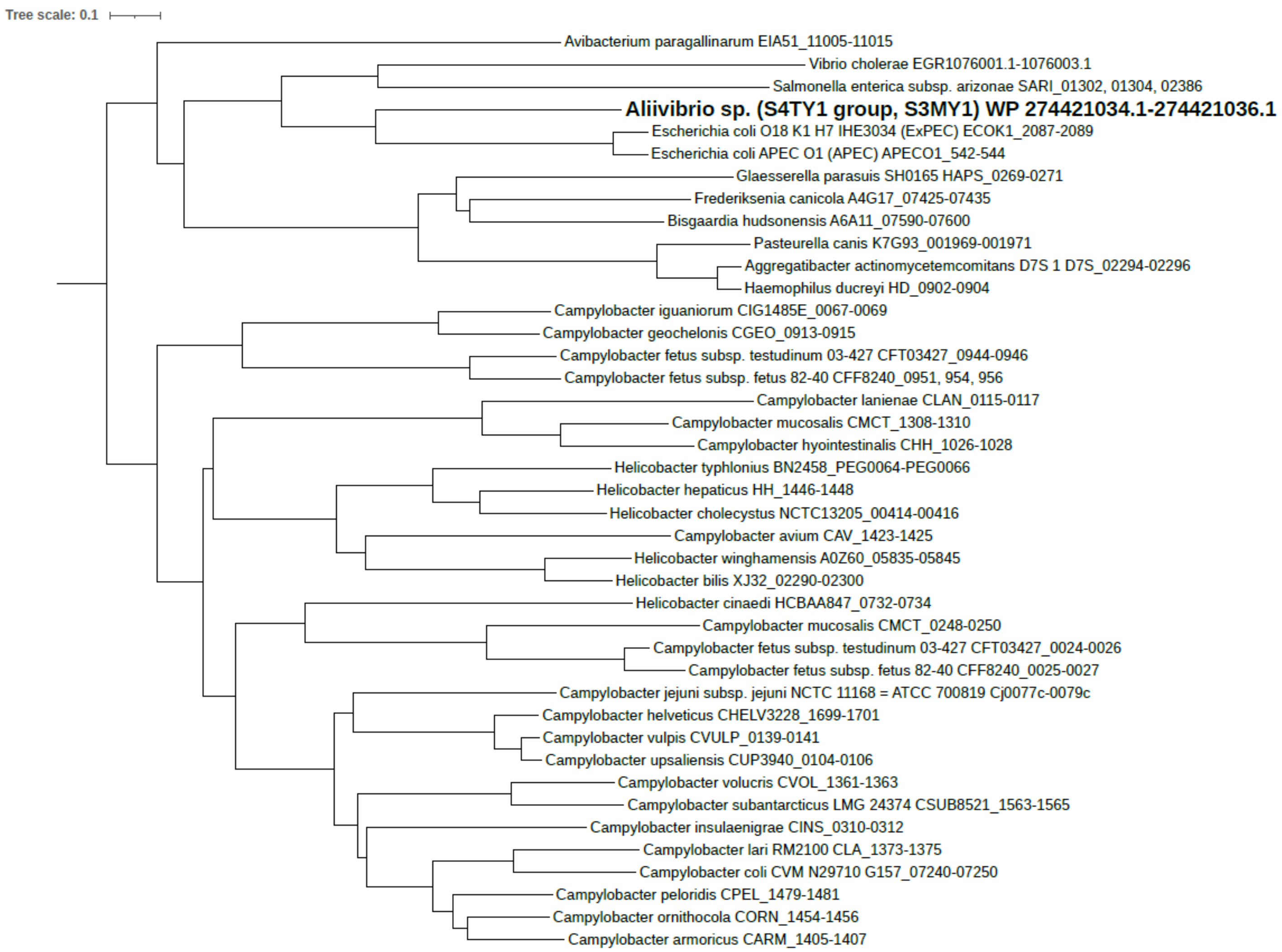


**Figure S6.** Group average-based tree based on concatenated cytolethal toxin subunit CdtA, CdtB and CdtC protein sequences. Only bacteria that included an intact operon are shown (most taxa shown are annotated in detail in the KEGG database). The CdtABC homologs present in *Aliivibrio* strain S3MY1 and strains of the S4TY1 group had identical amino acid sequences. Closest matches (55% identity overall) was to proteins of pathogenic *E. coli* strains (ExPEC – extraintestinal pathogenic E. coli, APEC – avian pathogenic E. coli). The Cdt sequences from a human faecal *Vibrio cholerae* isolate (no. 633012) deposited into NCBI by the England Public Health in 2020 (PM Ashton and colleagues) was more distantly related but in the same cluster. The expression, function and impact of Cdt in Atlantic salmon is currently unknown. Nor is it know the frequency of *cdt* genes in *Aliivibrio* or other fish-associated bacteria.
